# State modulation in spatial networks with three interneuron subtypes

**DOI:** 10.1101/2024.08.23.609417

**Authors:** Madeline M. Parker, Jonathan E. Rubin, Chengcheng Huang

## Abstract

Several inhibitory interneuron subtypes have been identified as critical in regulating sensory responses. However, the specific contribution of each interneuron subtype remains uncertain. In this work, we explore the contributions of cell-type specific activity and synaptic connections to dynamics of a spatially organized spiking neuron network. We find that the firing rates of the somatostatin (SOM) interneurons align closely with the level of network synchrony irrespective of the target of modulatory input. Further analysis reveals that inhibition from SOM to parvalbumin (PV) interneurons must be limited to allow gradual transitions from asynchrony to synchrony and that the strength of recurrent excitation onto SOM neurons determines the level of synchrony achievable in the network. Our results are consistent with recent experimental findings on cell-type specific manipulations. Overall, our results highlight common dynamic regimes achieved across modulations of different cell populations and identify SOM cells as the main driver of network synchrony.

## Introduction

As animals navigate the environment, their nervous systems process and react to an ongoing bombardment of sensory information. Internal factors such as motivation, attention, expectations, and arousal strongly impact animals’ perception, behavior and decision-making (*1–4*). Inhibitory neurons play an essential role in modulating the information processing and communication in cerebral cortex by tuning cortical oscillations, regulating the time window in which external inputs elicit cortical responses, and modifying the response gain of their excitatory counterparts (*5–7*). Inhibitory neurons, however, cannot be considered as a homogeneous population, but instead exhibit differences in morphology, connectivity, and biophysical properties (*8, 9*). Differences in molecular markers distinguish three non-overlapping inhibitory interneuron subtypes: parvalbumin (PV), somatostatin (SOM), and vasoactive intestinal peptide (VIP) expressing neurons. These interneuron subtypes are differentially targeted by neuromodulators and cortical feedback projections (*9–11*), and are thought to be involved in the modulation of neural population responses by brain state. Arousal and locomotion state of an animal have been shown to exert diverse influences on the firing rates of interneuron subtypes (*12–14*) and to strongly impact the synchrony level of neural population responses (*15, 16*). However, the functional role of each interneuron subtype remains unclear.

Advancements in optogenetic techniques enable the use of cell-type-specific stimulation and suppression to study the causal contributions to circuit dynamics by each cell type. Prior work demonstrated diverse effects on cortical firing rates and oscillations elicited by manipulating different target cell classes within cortical microcircuits (*6, 10, 17–20*). Stimulating PV neurons periodically enhances the oscillatory power of the local field potential (LFP) over the gamma frequency range (*21, 22*). This is consistent with previous theories where PV neurons are instrumental in generating gamma oscillations, partly due to their strong reciprocal connections with the excitatory neurons (*23*). However, recent work suggests that SOM neurons are involved in oscillations in the low gamma/beta frequency range (20-40 Hz), while suppressing PV neurons increases the spectral power of the LFP overall (*18, 24*). Suppressing SOM neurons also reduces the coherence between distant neural ensembles (*24*), consistent with their broad integration of lateral excitatory inputs (*25*). Stimulating VIP neurons increases the response gain of excitatory neurons, presumably through the disinhibitory pathway via SOM neurons (*26*). Silencing VIP neurons reduces the sensitivity of excitatory neurons to stimulus context (*27*) and increases the detection threshold for small visual stimuli (*28*). Despite the proliferative experimental findings, the network mechanisms underlying the observed changes in neural activity remain elusive, due to the intrinsic nonlinearity of the highly recurrently connected networks to which all of these cell types belong. Manipulation of one cell type leads to changes in the activity of the other cell types; however, experimenters typically observe the activity of all neurons indiscriminately or label one cell type at a time (but see (*29, 30*)). Therefore, computational models are needed to parse out the interactions between excitatory neurons and the three interneuron subtypes.

Previous models that incorporate multiple interneuron subtypes mostly focus on modulations of firing rates and do not consider impacts on network synchrony or correlations in neural activity (*31–35*). Some models have suggested that PV and SOM neurons contribute to oscillations of different frequencies (*36, 37*). However, these models are small networks or rate models and do not consider the spatial dependence of synaptic connections. In this work, we studied state modulation in spatially structured spiking neuron networks including multiple interneuron subtypes. Our past work has shown that such models can reproduce the irregular and weakly correlated neural population activity commonly observed in cortical recordings (*38*). We applied modulatory input to neurons of each cell type and analyzed the resulting changes in firing rates and network synchrony. We found that the pattern of activity changes resulting from activation of excitatory (E) or PV neurons is distinct from that due to activation of SOM or VIP neurons. Strikingly, SOM firing rates closely aligned with levels of network synchrony across all modulation cases. We further identified that stronger SOM→E than SOM→PV inhibition is important for maintaining a weakly synchronized dynamical regime, and that the interaction between E and SOM neurons is essential for enhancing network synchrony. Our work emphasizes the uniquely critical role of SOM neurons in regulating the dynamical state of cortical networks.

## Results

We developed a spatially-extended network model that includes one E population and three distinct inhibitory interneuron populations: PV, SOM, and VIP. Each neuron is modeled as a spiking exponential integrate-and-fire (EIF) unit (*39*). The synaptic connection patterns among the four neuron populations are constrained by anatomical and physiological data from mouse visual cortex (Fig. 1A) (*40*,*41*). In particular, we assume there are no reciprocal connections among SOM neurons or among VIP neurons; VIP neurons mainly inhibit SOM neurons, in what is believed to be an important disinhibitory pathway (*26*); and feedforward inputs only target E and PV neurons (*25*). Neurons are randomly distributed on a two-dimensional plane (1×1 mm^2^) and synaptic connection probability between neurons decays with distance (Fig. 1B; Equation 5). The spatial structure of the network allows for rich spatiotemporal activity patterns, such as propagating waves and spatiotemporal chaos, with population statistics consistent with cortical recordings (Fig. 1C, S14; Ref (*38, 42*)). Connections to and from the SOM cells have a larger spatial footprint compared to other connections, which is thought to be involved in surround suppression in visual cortex (*25*,*43*). The synaptic timescales of inhibitory connections from SOM and VIP neurons are slower than that of connections from PV neurons, which is in turn slower than that of excitatory connections, constrained by physiological data from mouse visual cortex (*44*). The network has a total of 50,000 neurons comprising 40,000 E, 4,000 PV, 4,000 SOM, and 2,000 VIP neurons, with the population size ratios following anatomical data from mouse cortex (*45*).

**Figure 1:**
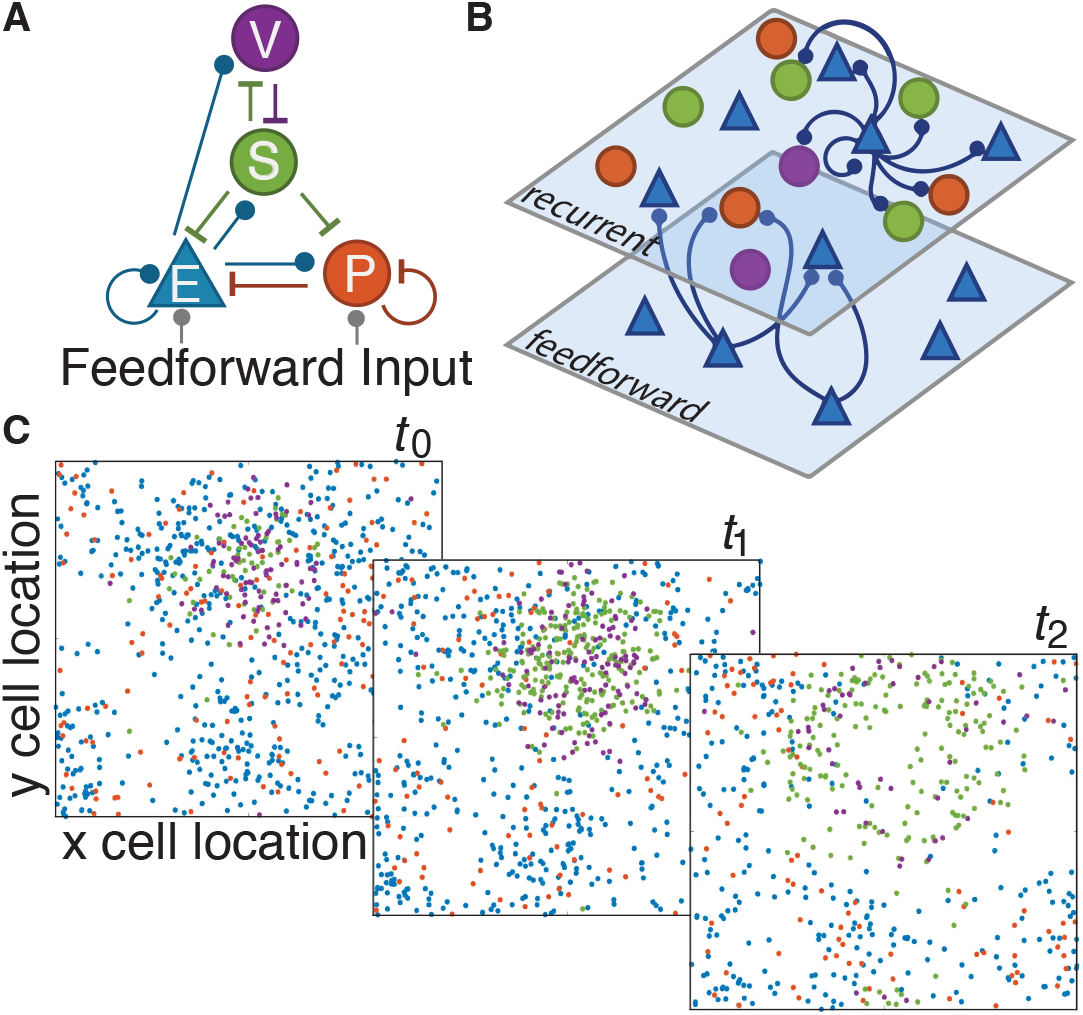
General model scheme and example dynamics. (A) Default network circuit diagram shows excitatory connections in blue (lines with circles) and inhibitory connections (T-lines) in other population-specific colors. (B) The model comprises one recurrent layer with one excitatory population and three inhibitory populations connected as in (A) and a feedforward layer, modeled as independent Poisson units, that provides excitatory input to E and PV neurons. Connection probability decreases with pairwise distance, as is illustrated schematically for E cells here. (C) Three consecutive spike raster snapshots, where a dot with a cell-type-specific color indicates that the neuron at spatial position (*x, y)* fired within 1 ms of the time stamp. In this example, local activity of E neurons (*t*_0_) recruits more activity of SOM neurons at a later time point (*t*_1_), which in turn suppresses the activity of all other neuron populations (*t*_2_).

### Network transitions through three dynamical states under variation of cell-type specific inputs

To begin our investigation, we apply a static input to one targeted population in the model. The static input remains constant in time and is applied equally to all the receiving neurons in the target population. In addition to the static input, E and PV neurons also receive feedforward inputs, modeled as Poisson processes, and all neurons receive recurrent synaptic inputs from other neurons (Fig. 1A, B; see Methods). In distinct trials, the static input is varied from -1 to 1 to examine the effects on network activity. We repeat this protocl with each of the four populations as the target population for the static inputs (Fig. 2, 3). We find that the network exhibits three typical dynamical states as the static input value is varied. We term these three states as subcircuit asynchronous (SA), weakly synchronous (WS) and strongly synchronous (SS) states (Fig. 2). We first define each state and then examine the effects of input modulation for each target population.

**Figure 2:**
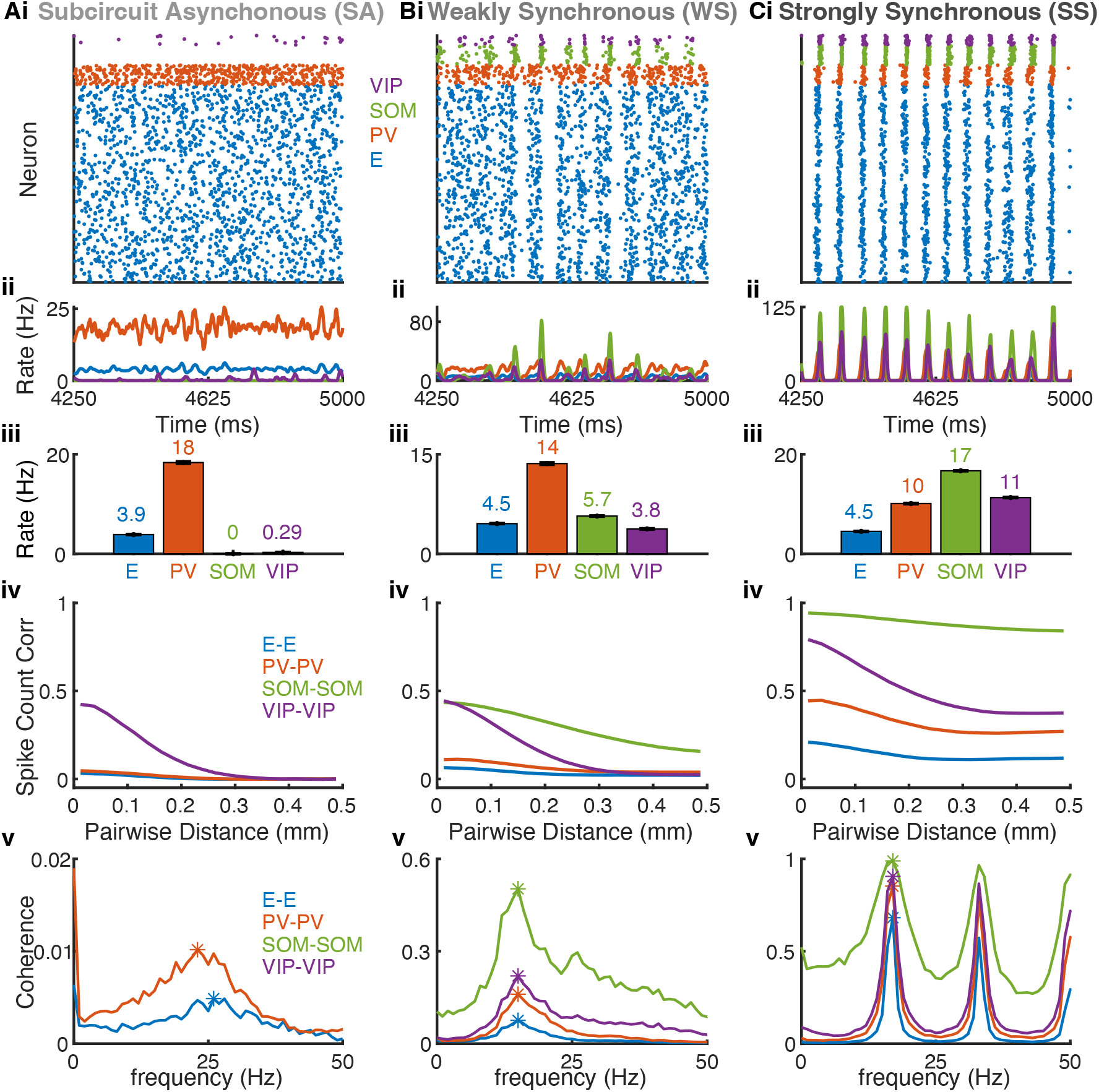
Three representative network states. SA state (Ai-v), WS state (Bi-v) and SS state (Ci-v). Row (i): Spike raster of a subsample of each of the four populations: 400 E (blue), 40 PV (red), 40 SOM (green), and 20 VIP (purple) neurons. The number of neurons of each neuron population shown in the rasters is proportional to the population size. Row (ii): Population-averaged firing rates over the same time course as the spike rasters in row (i). Row (iii): Mean firing rates averaged over neurons and over time for each population. The number on top of each bar is the value of the mean firing rate. Error bars are standard error of mean (SEM). Row (iv): Average spike count correlations (see Methods) as a function of distance for neuron pairs within each population. The networks spans a 1×1 mm^2^ square. Row (v): Average pairwise coherence of spike trains (see Methods) versus frequency for neuron pairs within each population. The asterisks mark the maximum coherence over non-zero frequencies. Note the different y-axis scales across panels. The presented cases correspond to input to PV with static input = 0.6 (column A), static input = 0.0 (column B), and static input = -0.6 (column C).

**Figure 3:**
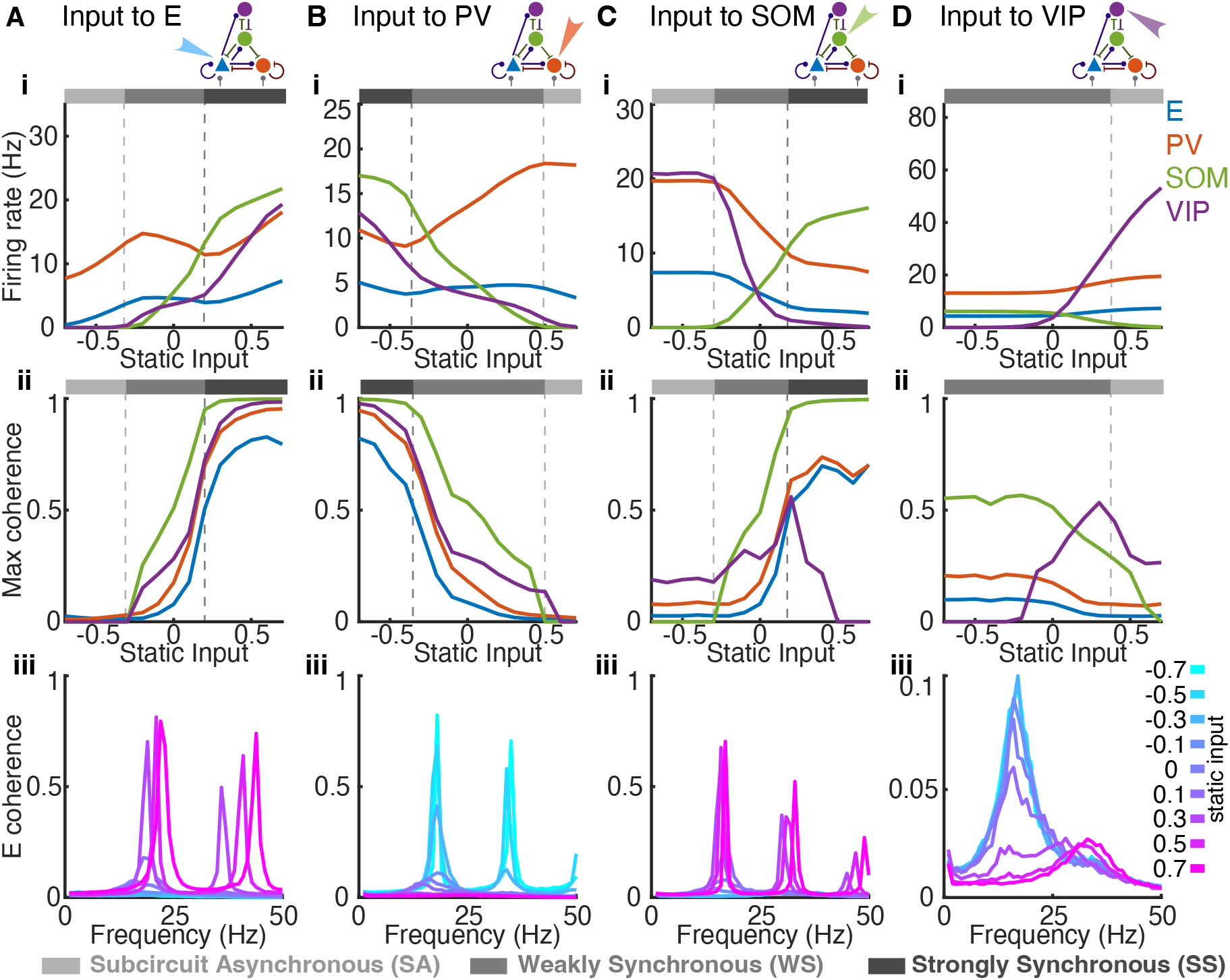
Cell type specific inputs change population firing rates and coherence in distinct ways. Static input is applied to all neurons in one of the four populations: (A) E, (B) PV, (C) SOM or (D) VIP neurons. Row (i): Average population firing rates of each population as a function of static input value. Grey-scale bars above each plot represent network activity state at the corresponding input value (SA: light grey; WS: moderate grey; SS: dark grey). Note the differences in vertical axis scales across panels. Row (ii): Maximum coherence in each population as a function of static input value. Row (iii): E population coherence as a function of frequency for several static input values. Note the distinct vertical axis scale in panel D(iii).

We define activity states based on measurements of average firing rates and spike train coherence (see Methods). Specifically, the SA state is defined as when average firing rate of SOM neurons is less than 1 Hz and the maximum coherence of E neurons is less than 0.1. The WS state is when the maximum coherence of E neurons is between 0.1 and 0.5 and when average firing rate of SOM neurons is larger than 1 Hz. The SS state is when the maximum coherence of E neuron is larger than 0.5.

In the SA state (Fig. 2Ai-iv; Supplemental video 1), the network behaves essentially as a two population subcircuit composed of interacting E and PV neurons, while SOM and VIP activity is nearly, if not completely, silent. The E population is the only excitatory source of input to SOM and VIP. In the SA state, E neurons are unable to consistently drive SOM and VIP neurons over their respective spiking thresholds (Fig. 2Aii-iii). E neurons exhibit little synchronization or organized activity, as indicated by the near-zero levels of average spike count correlations between E neuron pairs (Fig. 2Aiv). The average spike train coherence among E or among PV neurons is also low with a peak above 25 Hz (Fig. 2Av).

Within the WS state, all four populations actively fire (Fig. 2Bi-iv). PV neurons exhibit the highest firing rates, with the other three populations moderately active (Fig. 2Bi-iii). The spiking activity of E and PV neurons is largely asynchronous, interspersed with brief coordinated periods of silence (Fig. 2Bi; Supplemental video 2). The silent periods in E and PV are preceded by synchronous bouts of rapid firing in SOM and VIP neurons (Fig. 2Bi,Bii). The spike count correlations and coherence of SOM-SOM and VIP-VIP neuron pairs are larger than those of E-E and PV-PV neuron pairs (Fig. 2Biv-v), consistent with experimental observations in mouse cortex (*44, 46*). The correlation between SOM neuron pairs persists over larger distances than those of other populations, due to the larger spatial footprints of SOM neuron connections, which is also consistent with cortical recordings (*46*).

The SS state exhibits highly synchronized and oscillatory activity in all populations (Fig. 2Ci-iv). Patterns of firing initially begin with a low number of E and PV spikes, which recruit many more E and PV neurons to fire, thereby activating a large portion of SOM and VIP neurons (Supplemental video 3). The elevated firing rates of all three inhibitory populations (Fig. 2Cii-iii) supply a substantial amount of inhibitory current, ultimately silencing all neurons until enough feedforward input accumulates to excite E and PV neurons and to cause the cycle to repeat (Fig. 2Ci). Pairwise spike count correlations are relatively large within each population and only slightly decrease with distance (Fig. 2Civ). The average coherence of each neuron population shows a dominant peak close to 1 at around 20 Hz (Fig. 2Cv). Since spike count correlations depend on the choice of time window for calculating spike counts, the correlation value can be misleadingly low when the time window coincides with the multiples of the oscillation period. For this reason, we hereafter use the maximum coherence to measure the level of network synchrony.

Comparing the effects of varying a static input current applied to different neuron populations reveals that external inputs to different targets modulate population dynamics across similar states. Specifically, we see that activating E neurons increases coherence in all populations (Fig. 3A). As input to E increases, network activity transitions from the SA to the WS to the SS state. The transition is marked by non-monotonic changes in firing rates in E and PV populations (Fig. 3Ai). The firing rate of E neurons decreases with increasing static input drive in the WS state, which is counterintuitive. On the other hand, the firing rate of SOM neurons monotonically increases in the WS state, despite of the reduction in E firing, which is the sole source of excitatory inputs to SOM neurons.

To explain the changes in population rates with input drive, we estimated the rate-current transfer functions of the EIF neuron model of each cell type, with colored noise of the same timescale as the excitatory synapses (fig. S1). The transfer functions depend on the mean and the variance of the total input current each neuron receives, which we measure from network simulations (fig. S1, S2iii). The prediction of population rates using the rate-current transfer functions of single neurons matches those from network simulations closely when the network is in the SA or WS state, but differs when the network is in the SS state (fig. S2iii). The failure of this prediction in the SS state is due to the strong oscillation and non-Gaussianity in synaptic inputs from the network. In general, firing rate increases monotonically with mean current and increases with the variance of current when mean current is below or just above threshold (fig. S1D), consistent with previous results with white noise input (*47*). In the transition from the WS to the SS state, the variance of input current rises rapidly as the static input to E neurons increases, which drives the SOM neurons to a higher rate despite the reduction in mean current (fig. S2Ai-ii). When fixing the current variance at the value from the SA state (input = -1), the predicted SOM rates remained zero until input is above 0.5 (fig. S2Aiv, circles). The large increase in SOM firing rate in turn results in enhanced inhibition from SOM to E neurons (fig. S3Ai), which leads to a reduction in E firing rate. In contrast, E firing rate increases monotonically with input applied to E neurons when solving the population rates self-consistently with fixed rate-current transfer functions, assuming that the current variances are independent of static input values (fig. S2Av). This suggests that the changes in current variance qualitatively change the dependence of population rates on input drive.

When increasing the external input to PV neurons, we observe a reverse order of state transitions compared to the case with input to E neurons (Fig. 3B). Activating PV neurons decreases coherence in population spiking, moving the network from the SS to the WS to the SA state. The response gain of the PV neurons increases by several folds as the external input to PV neurons increases (fig. S4Bii), which means that the PV neurons become more effective at suppressing the rate fluctuations of the E neurons. In addition, in the SS state, larger external input to the PV neurons makes them fire earlier in each oscillation cycle and reduces the magnitude of E firing rate (fig. S5). The firing rates of E and PV neurons again exhibit non-monotonic changes, as in the case with input to E neurons (Fig. 3A). In the WS state, driving PV leads to small increases in E firing rate because of the reduction in the inhibition from SOM neurons (fig. S3Bi). The firing rates of SOM drop despite the presence of increases in mean excitation due to the large reduction in the variance of input current (fig. S2Bi-ii). When fixing the current variance at the value from the SS state (input = -1) and predicting rate based on mean current values from network simulations, the predicted SOM firing rates remained high and increased with static input levels in the WS state (fig. S2Biv).

Driving SOM neurons increases population coherence and moves the network from the SA to the WS to the SS state, similar to the effects observed when driving E neurons (Fig. 3C). However, the firing rates of E and PV neurons monotonically decrease as SOM neurons become more active (Fig. 3Ci), in contrast to the non-monotonic changes that result when driving E or PV neurons (Fig. 3Ai,Bi). VIP neurons become suppressed when SOM neurons are sufficiently activated due to inhibition from SOM to VIP (Fig. 3Ci). In contrast, VIP and SOM firing rates co-vary in the same direction when input is applied to E or PV neurons (Fig. 3Ai,Bi).

Lastly, varying the external input to VIP neurons yields similar changes to those arising with PV input variations, but is unable to induce all three of the network states that we have identified (Fig. 3D). When input to VIP is strong, inhibition from VIP to SOM shuts down SOM activity and firing rates of E and PV neurons increase slightly due to disinhibition. With SOM silenced and VIP having no synaptic connections to PV and E neurons, the network behaves asynchronously and effectively like a two population E-PV subcircuit, thus adopting the SA state. When input to VIP neurons is reduced, the drop in inhibition from VIP to SOM means that SOM starts to fire and VIP firing decreases. The network transitions from the SA to the WS state, and stays in the WS state once VIP neurons are fully suppressed (Fig. 3Di, ii). Therefore, modulating VIP neurons does not lead the system to the pathological SS state, which makes the VIP neurons a ideal candidate for moderate state modulations. In addition, we observe that the frequency of peak coherence within the E population transitions (Fig. 3Diii, S6D) as a result of changes in static input to VIP neurons. Activating VIP results in peak frequencies occurring at around 30 Hz, but as VIP reduces its activity due to reduced input and SOM begins to fire, the peak frequency shifts to approximately 15 Hz, with higher peak levels of E coherence.

In all input cases, we observe similar network states for a given input in multiple simulation runs with random initial conditions. We also did not observe bistability when comparing network activity with gradually changing (increasing or decreasing) input (fig. S11A,B). Based on the absence of hysteresis effects, we infer that the transition from the SA to the WS state likely occurs through a supercritical Hopf bifurcation.

### Firing rates of SOM neurons co-vary with network synchrony

To directly compare how firing rates and network synchrony change together as input to each neuron population varies, we summarize the results of four input cases from the previous section with phase plots of the maximum coherence of the E population versus the firing rate of each neuron population (Fig. 4). On these phase plots, each trajectory corresponds to a path of network state transitions as input to a specific neuron population varies. The arrows represent the directions of transition as input increases value. We use the maximum coherence of the E population to represent the overall network synchrony level for two reasons: first, E neurons make up the majority of the total neuron population (80%) and are recorded most commonly in experimental research and secondly, the coherence of all four populations tend to vary together, other than some exceptional results in VIP neurons (Fig. 3A-Dii).

**Figure 4:**
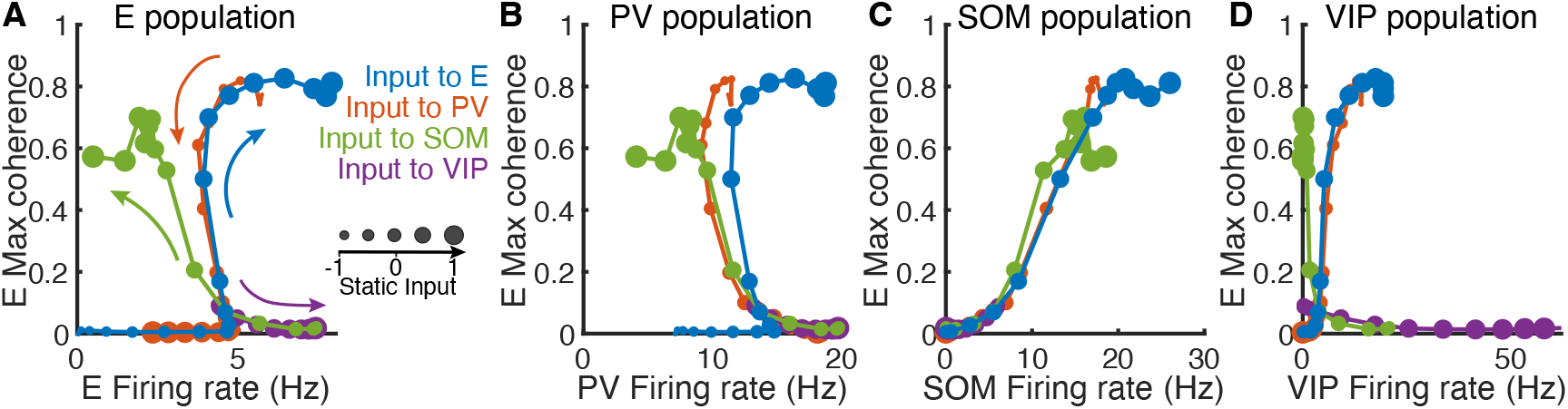
Modulation patterns of population firing rates and E population coherence. Levels of external input are indicated by individual circular markers, where decreasing marker size signifies decreased static input (i.e., progressing from activating to suppressing the target). Colors in each panel indicate which population receives the varying input and colored arrows (A) indicate the direction of increasing input (following the direction of increasing marker size along a single colored curve). Each panel depicts E coherence versus the population-averaged firing rate of one neuron population: (A) E, (B) PV, (C) SOM, and (D) VIP.

When plotting E population coherence versus E population firing rates, we find that for the cases of input to E or PV neurons, the network evolves along a common path, with opposite directions of traversal resulting from similar changes in static input levels (Fig. 4A). Similarly, we obtain a common path for the cases of input to SOM or VIP neurons, but this common path differs from that observed with input to E or PV neurons. Previously we found that input to VIP never resulted in SS activity (Fig. 4Dii), which explains why the VIP curve (purple) ends at a relatively low coherence value. Within the common path shared by E and PV stimulation, there exist three regimes: a lower branch where coherence is low (∼ 0) and input changes only affect firing rate (the changing markers on the *x*-axis), an upper branch where coherence remains high (*>* 0.5) over a range of high firing rates, and a transition between the low and high coherence plateaus, across which coherence changes substantially while firing rates remain relatively unchanged. These regimes align with the network activity: the lower branch is the SA state, the upper branch is the SS state, and the transition is the WS state. What is especially remarkable is the precise overlap of pairs of paths, along with the alignment of all paths during the transition region (i.e. the WS state), which suggests that the network structure strongly constrains network dynamics. The modulation patterns in the full network are distinct from those in the isolated E-PV subcircuit, where firing rate and coherence levels tend to vary in the same direction and monotonically as input level varies (fig. S7).

Similarly, curves of E coherence versus PV firing rate overlap mostly for E and PV input cases, as do the curves for SOM and VIP input cases (Fig. 4B). When comparing the E population coherence and SOM population rates (Fig. 4C), however, the paths corresponding to application of static input to all four target populations largely overlap, no longer showing a distinction across input targets (aside from the direction of modulation across states as indicated by changes in marker sizes). Lastly, VIP firing rates compared to E coherence for all input cases (Fig. 4D) features the dichotomy of trajectories generated by inputs to E and PV versus paths from inputs to SOM and VIP (as also observed in Fig. 4A, B).

Overall, we see that applying excitatory input to E or SOM neurons or inhibitory input to PV or VIP neurons tends to increase coherence, although this change is accompanied by distinct changes in firing rates for most cell populations. Comparing SOM population rates with the coherence within the E population, however, reveals that the two quantities increase together, in a stereotyped way, in all input cases (Fig. 4C). This consistency between E coherence and SOM activity across all input targets leads us to postulate that SOM activity plays a central role in dictating the level of network synchrony.

### Strong SOM inhibition to PV drives synchrony

We next investigate how synaptic connection strengths in the network shape the modulation patterns of network states induced by cell-type specific inputs. Building on our prior observation of the alignment of SOM firing rate with network synchrony, we focus on the strengths of connections projecting onto or from SOM neurons, specifically SOM→E, SOM→PV, and E→SOM (next section) synapses. Since the influence of VIP’s inhibitory outputs is restricted to SOM neurons, varying the connection strengths between VIP and SOM neurons has little effect on the input-induced transition patterns (fig. S8 and S9).

We find that SOM→E connections are important for generating the non-monotonic changes in E and PV firing rates along the transition paths induced by varying input to E or PV neurons (Fig. 3Ai, Bi, and 4A). When we eliminate SOM→PV connections (i.e., *J*_*SOM*→*PV*_ = 0), modulation patterns in network activity states (Fig. 5Ai, Bi) remain qualitatively the same as in the network’s default setting (Fig. 3Bi, and 4Ai, Bi; more combinations of SOM→E and SOM→PV connection strengths are in fig. S10). Stronger SOM→E connections lead to a larger range of firing rates over the transition from the SA to the SS state through the middle branch, corresponding to the WS state, where rate and coherence vary in opposite directions (Fig. 5Ai). The changes in state in response to external input variations are gradual in networks with different SOM→E connection strengths (Fig. 5Ai, Bi). This points to a degree of resilience in the network’s responsiveness to external input in the absence of SOM inhibition to PV.

**Figure 5:**
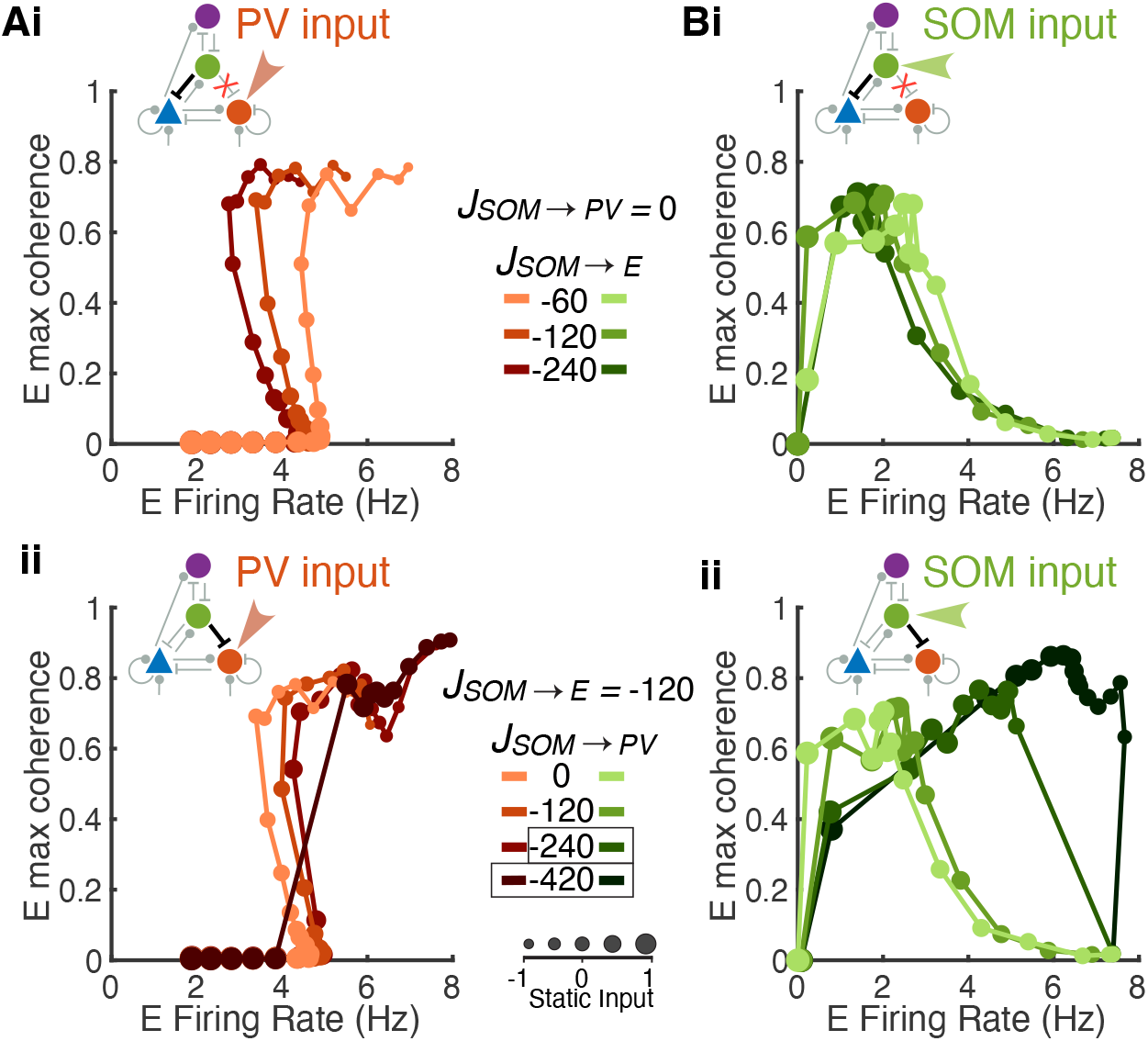
Relative strengths of SOM →E and SOM →PV connections shape modulation patterns of network state. Static input is applied to either PV neurons (Ai, ii) or SOM neurons (Bi, ii). Network state at each input level is represented by E firing rate and E maximum coherence (with the same convention as in Fig. 4A). Increasing marker sizes correspond to increasing static input to the target population. (i) SOM inhibition to PV is removed (*J*_*SOM*→ *PV*_ = 0) and increases of *J*_*SOM*→ *E*_ correspond to darker curves. (ii) SOM inhibition to E is fixed (*J*_*SOM* →*E*_ = −120) and increases of *J*_*SOM* →*PV*_ correspond to darker curves. Default values of connections strengths are *J*_*SOM* →*E*_ =− 120 and *J*_*SOM* →*PV*_ = −60. Boxed connections weights highlight networks with abrupt transitions in the modulation patterns of network activity.

This resilience is disrupted when the synaptic strength of SOM→PV dominates the synaptic strength of SOM→E, which results in an increased sensitivity of the network to changes in external input. Specifically, we consistently observe that when SOM inhibition to PV is sufficiently large compared to SOM inhibition to E, more pronounced and abrupt transitions from the SA to the SS state occur (e.g., the cases highlighted with black boxes in Fig. 5Aii with *J*_*SOM*→*PV*_ = −420 and in Fig. 5Bii with *J*_*SOM*→*PV*_ = −240, −420; fig. S10). That is, dominance of SOM→PV inhibition over SOM→E inhibition increases network sensitivity to input and reduces or eliminates the range of input levels that result in the transitional activity dynamics, the WS state. Indeed, in the transition through the WS state, as SOM firing intensifies (Fig. 3i), the inhibition from SOM to E and PV neurons will tend to reduce their firing rates. Yet, the drop in PV firing can disinhibit E. If this disinhibitory effect is dominant due to sufficiently strong *J*_*SOM*→*PV*_ , then E firing can increase rather than decreasing, resulting in a rapid transition through or elimination of the WS state. In this case, the firing rate and maximum coherence of E neurons tend to vary in the same direction (Fig. 5ii, fig. S10). Comparing results from increasing and decreasing incremental changes in input levels, we observe that the abrupt transition between SA and SS states happens at different input values depending on the direction of change (fig. S11). This hysteresis effect suggests that stronger SOM inhibition to PV neurons changes the criticality of the Hopf bifurcation at which SA stability is lost, from supercritical to subcritical.

These results imply that stronger inhibition from SOM→E neurons than that from SOM→PV neurons is necessary to observe activity consistent with the WS state and underscores the pivotal influence of SOM inhibition on the network’s dynamical transitions.

### Dynamic interactions between E and SOM neurons are necessary for SOM-induced network synchrony

In this section, we investigate the impacts of E→SOM connections on SOM-induced network synchrony. What might drive the high coherence among SOM neurons and the rest of the newtork? Since SOM neurons do not connect to other SOM neurons and do not receive feedforward input, the high correlation among SOM neurons is driven by the recurrent input they receive from within the network. There are only two sources of recurrent inputs to SOM neurons, the excitation from E neurons and the inhibition from VIP neurons. To investigate the importance of E→SOM connections, we removed the E→SOM connections, and replaced this recurrent excitation with an external input that mimicked the statistics of the recurrent excitation.

First, we replaced recurrent excitation with colored noise that was independent for each SOM neuron. The colored noise was constructed as an Ornstein–Uhlenbeck (OU-) process that had equal mean and variance to the excitatory currents SOM neurons received on average in a intact default network with no static input (referred to as *baseline*; mean = 0.65 and variance = 0.12). In this decoupled network, the coherence of the network remains low and decreases as we apply static input to SOM neurons, in addition to the noisy input, to increase their firing rates (Fig. 6A). This result is opposite to the large increase in coherence with SOM rate that we observed in the default network (Fig. 4A,C). The firing rate of E neurons is also suppressed much more abruptly compared to that in the default network as we increase input to SOM neurons (fig. S12 compared to Fig. 3C). This suggests that without E→SOM connections, SOM activity tends to reduce network synchrony mainly by reducing the E firing rate. The inhibition from VIP alone is not able to correlate SOM neurons. Consistently, varying VIP→SOM connections has little effect on network coherence (fig. S9).

**Figure 6:**
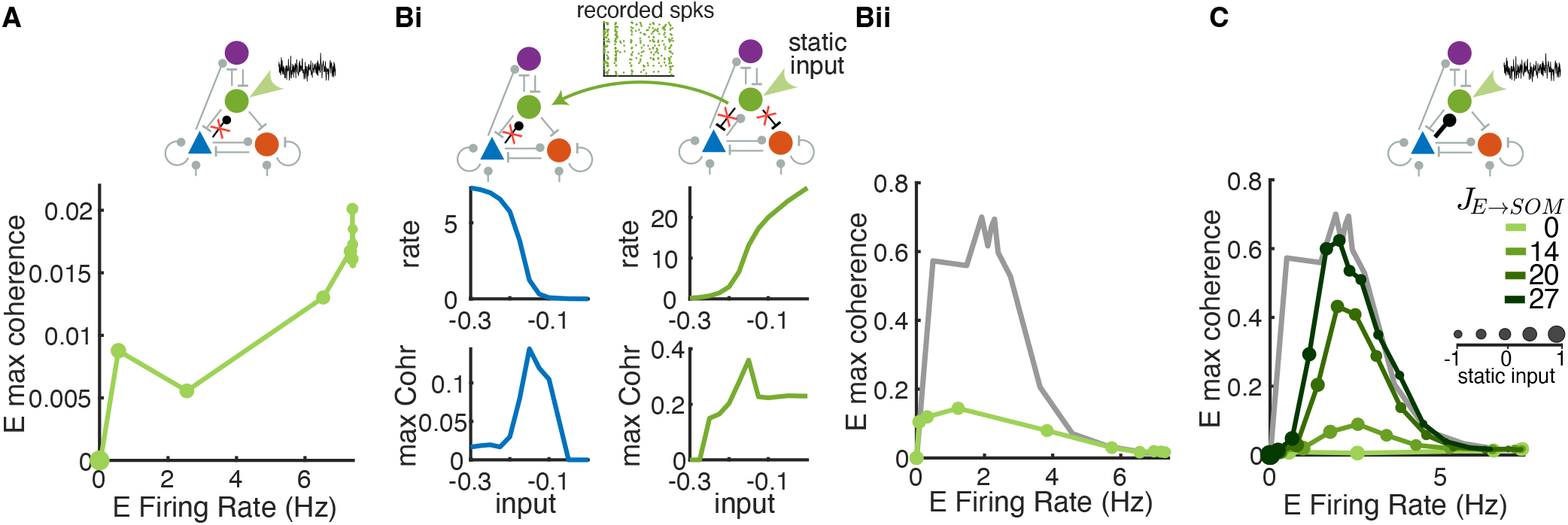
E→SOM connections are critical for SOM-induced network synchrony. (A) Removal of E→ SOM connections eliminates coherence, despite the presence of stochastic input (OU process, see Methods) with mean and variance matched those of the excitatory currents to SOM neurons in the intact default network with no static input (mean = 0.65 and variance = 0.12). Larger dots correspond to stronger static inputs to SOM. (Bi-ii) E and SOM firing properties in networks where they are dynamically uncoupled, but SOM neurons receive and provide realistically correlated inputs and outputs, respectively. (Bi) Left column: Firing rate (top) and maximum coherence (bottom) of E neurons in networks with no E →SOM connection and where SOM spikes were replaced with those recorded from the network on the right. Right column: Firing rate (top) and maximum coherence (bottom) of SOM neurons in networks with intact E→ SOM but no SOM →E and SOM →PV connections. Static input was applied to SOM neurons in the network on the right. (Bii) Modulation pattern of the firing rate and maximum coherence of E neurons from the network in Bi left (green) and from the default network (grey) with changes in static input to SOM neurons. (C) Increasing E →SOM synaptic strength increases the maximum coherence that can be achieved by varying static input to SOM neurons. SOM neurons receive the same OU noise as in panel A as *J*_*E* →*SOM*_ values are varied. Hence the lightest green curve (*J*_*E* →*SOM*_=0) is the same as that in panel A. As in panel Bii, the grey curve shows coherence for the default network with static input for comparison (same data as in Fig. 4A green curve).

Next, we consider the possibility that correlated excitatory inputs are able to synchronize SOM neurons, which in turn synchronize the network as a whole. Because each SOM neuron receives input from a large number of E neurons (∼1200 connections), very weak correlation in E spike trains can result in large correlation in the pooled excitatory current, as has been demonstrated theoretically (*48*). The correlated excitatory current to SOM neurons cannot be dynamically canceled by inhibition due to the lack of inhibitory connections among SOM neurons, which is distinct from the E-PV subcircuit where a balance of excitation and inhibition can be dynamically achieved (*49, 50*). Therefore, excitatory input alone is able to drive correlated activity in SOM neurons.

To demonstrate that correlated firing of SOM neurons can be driven by excitatory input alone, we record SOM spike trains from networks where SOM neurons receive excitation from E neurons but do not provide feedback inhibition to E and PV (Fig. 6Bi, right column). Static input is applied to SOM neurons to modulate their firing rate. We then replay the recorded SOM spikes in networks where we remove E→SOM connections but allow SOM neurons to impact the rest of the network (Fig. 6Bi, left column). In this way, E and SOM neurons are dynamically uncoupled, but SOM neurons receive realistic correlated excitation instead of simplified independent noise as in Fig. 6A. We find that as input to SOM neurons increases, firing rate of SOM rises rapidly and their coherence level reaches to about 0.3 (Fig. 6Bi, right column). This is consistent with the previous theoretical result that correlation between uncoupled neurons increases with firing rates (*51*). The increased coherence in SOM spiking activity in turn induces synchrony among E neurons until E neurons are fully suppressed by the inhibition from SOM (Fig. 6Bi, left column). Therefore, the correlated excitatory current to SOM neurons is able to drive the network into a weak synchrony regime (coherence around 0.15), but the peak coherence is much lower than that in the default network with E→SOM connections (Fig. 6Bii).

Lastly, as we gradually restore E→ SOM connections (*J*_*E*→*SOM*_ *>* 0) to allow for dynamic interaction between E and SOM neurons, we observe a positive relationship between the increases in coherence and increases in connection strength (Fig. 6C). These results demonstrate that mimicking E→SOM input, using either colored noise with matched mean and variance (Fig. 6A) or recorded SOM spikes from a decoupled network (Fig. 6Bi-ii), is not sufficient to modulate activity through the three identified network states; rather, it is the dynamic interaction between E and SOM neurons that amplifies the weak correlation in the E-PV subcircuit and drives the network to strong synchrony.

### Impacts of the spatial and temporal scales of SOM connections on network synchrony

Next, we investigate how network synchrony depends on the spatial and temporal scales of synaptic connections in our network. In our default network, the widths of connections both from and to SOM neurons (i.e., E→SOM, VIP→SOM, SOM→E, SOM→PV, SOM→VIP connections) are twice the widths of all other connections, based on data from mouse cortex (*25, 43*). The broad connections of SOM neurons could contribute to the global synchrony induced by SOM activity.

We find that it is the width of E→SOM connections that critically determines the level of coherence observed in the network. When the width of E→SOM connections is equal to or narrower than the rest of the connections, the maximum coherence observed with modulation is largely reduced (Fig. 7Aii, fig. S13B). In contrast, narrowing other SOM connections (i.e., VIP→SOM, SOM→E, SOM→PV, SOM→VIP connections) while keeping E→SOM connections broad does not change the coherence level across modulation input values (Fig. 7Aiii, fig. S13C). Interestingly, narrowing all five SOM connection types (both from and to) further reduces the maximum coherence across input levels (Fig. 7Aiv, fig. S13D). Overall, the trend of modulations in network synchrony with modulatory input remains the same in networks with different SOM connection widths, but the magnitude of coherence is larger when E→SOM connections are broader.

**Figure 7:**
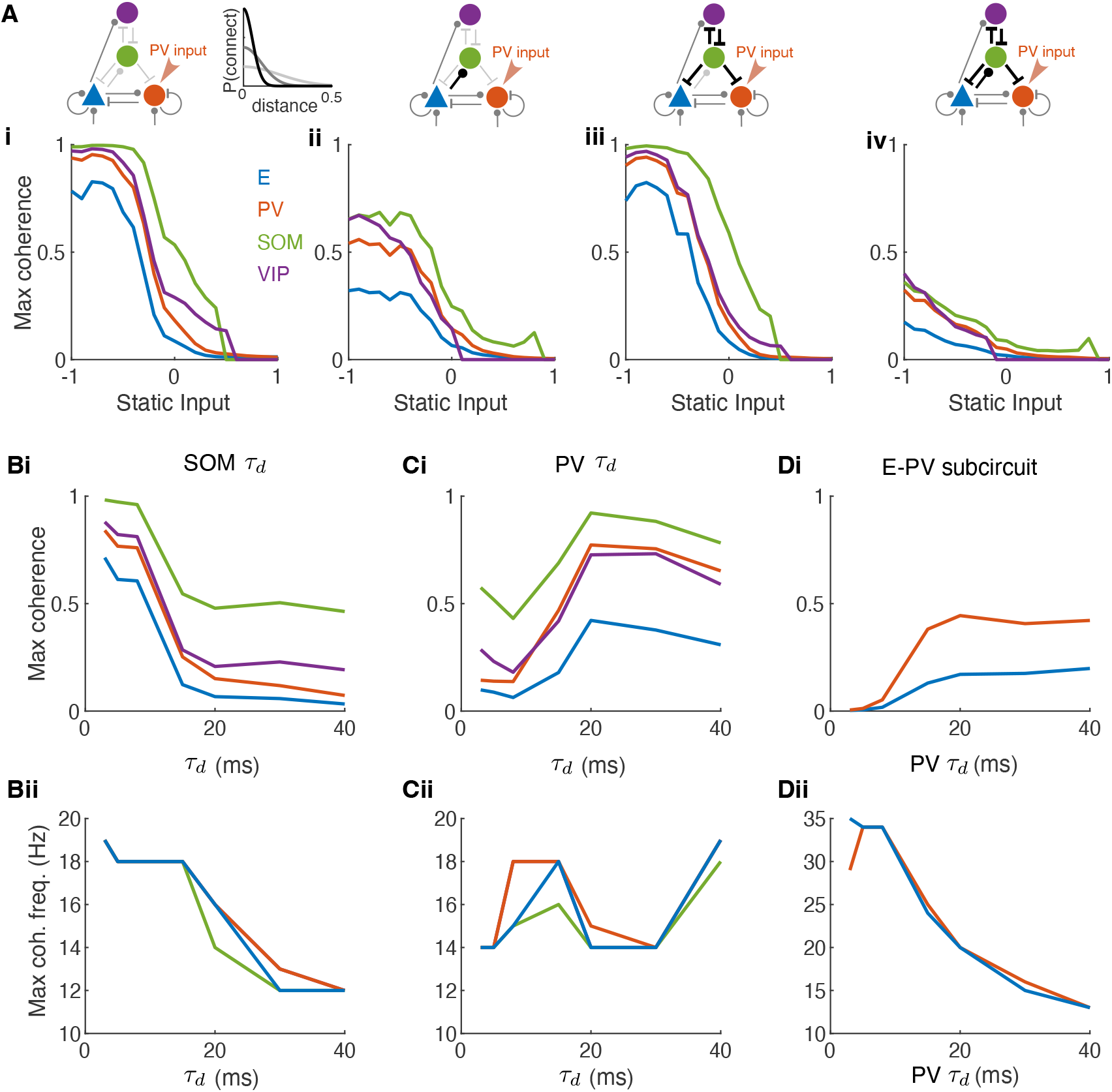
Impacts of the spatial and temporal scales of SOM connections on network synchrony. (A) Maximum coherence in each population as a function of static input value in networks with different spatial widths of SOM connections. Static input is applied to the PV neurons. Color of the connections in network schematic indicates the projection width *σ* (Equation 5). Light gray: *σ* = 0.2 mm; Medium gray: *σ* = 0.1 mm; Black: *σ* = 0.05 mm. (Ai), default network where connections to and from SOM neurons are broader (width = 0.2 mm) compared to other connections (width = 0.1 mm), same as Fig. 3Bii. (Aii), E →SOM connection width is narrowed to 0.05 mm while keeping all other connection widths the same as those in the default network. (Aiii), Other SOM connections, VIP→ SOM, SOM→ E, SOM →PV and SOM→ VIP connections, are narrowed to 0.05 mm width while keeping all other connection widths the same as those in the default network. (Aiv), all connections from and to SOM neurons are narrowed to 0.05 mm, while keeping all other connection widths the same as those in the default network. (B) Varying the decay time constant (*τ*_*d*_) of SOM synapses. (C) Varying the decay time constant of PV synapses. (D) Same as B in the E-PV subcircuit. Row (i): Maximum coherence in each population as a function of *τ*_*d*_. Row (ii): The frequency of maximum coherence in each population as a function of *τ*_*d*_. The static input was zero in B-D. *τ*_*d*_ is 8 ms and 20 ms for the PV and SOM synapses, respectively, in the default network in Fig. 3.

In addition, we find that the spatial structure in the network is important for gradual state transitions, consistent with the observations in our previous work (*38*). In networks with no spatial structure, meaning that the connection probability between two neurons does not depend on distance, we observe sharp transitions between SA and SS states as external input varies (fig. S14Ai-iv). Therefore, the spatial structure of the network contributes to maintaining a WS state over a range of input values.

Next, we analyze how synaptic timescales affect the synchrony level and the oscillation frequency in the network. We find that variations of the synaptic decay time constants of SOM 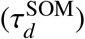 and 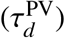 neurons have opposite impacts on network synchrony. Decreasing 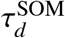 increases the maximum coherence in the network (Fig. 7Bi, fig. S15A). Since the integral of each synaptic current is normalized to be the same as we vary 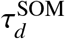 (Equation 3), a shorter 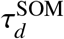 corresponds to a larger peak magnitude of each post-synaptic inhibition from SOM. Therefore, the SOM neurons with shorter 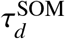 could more effectively override the E neurons and induce global oscillations in the network. The peak frequency of maximum coherence increases as 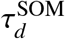 decreases, but plateaus at about 18 Hz for 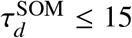 ms when the network is in the SS state (Fig. 7Bii). This suggests that 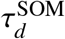 contributes to the oscillations in the beta frequency range (15-20 Hz) when the network is in the WS state. The oscillation frequency is relatively insensitive to 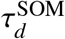 when the network is in the SS state.

In contrast, reducing 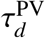 below 20 ms generally decreases the maximum coherence (Fig. 7Ci), consistent with our previous findings (*38*). This is because PV neurons with shorter 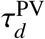 are more effective at suppressing E and other PV neurons whenever there is a rise in E population rate. The coherence of the E neurons remains relatively low as we vary 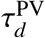 from 3 to 40 ms. The peak frequency of maximum coherence varies between 14 and 19 Hz and does not show consistent patterns (Fig. 7Cii, fig. S15B). To isolate the impacts of PV synaptic timescale without the interaction with SOM neurons, we analyzed the effects of 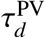 in the E-PV subcircuit (Fig. 7D). Reducing 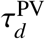 in the E-PV subcircuit monotonically decreases the maximum coherence of the E neurons and increases the peak frequency (Fig. 7Di-ii, fig. S15C). This demonstrates that the synaptic timescale of PV neurons contributes to the oscillations around 35 Hz that we observed in the SA state (Fig. 3Diii).

### Heterogeneous external inputs reduce SOM-induced network synchrony

In our previous set of results, adding noise to SOM neurons only slightly reduced the coherence of the network when E firing rate is small (Fig. 6C, compare dark green with grey curves). This observation suggests that the network can still readily transition into a highly coherent regime even in the presence of noisy inputs that vary in time. To investigate the impact of noise in the external input, we applied independent OU input, with equal mean and variance, to each neuron in the SOM population (see Methods). Increasing the variance of the OU input only weakly impacted the coherence of the network (Fig. 8A). We next compared this outcome with the results of applying a different type of applied noise, heterogeneous or quenched noise, which is constant in time but has heterogeneous strengths, sampled from a normal distribution of a given variance, across target neurons (see Methods). Implementing the OU input and the quenched input allows us to compare the role of time-varying noise versus spatially-varying noise in terms of influence on coherence.

**Figure 8:**
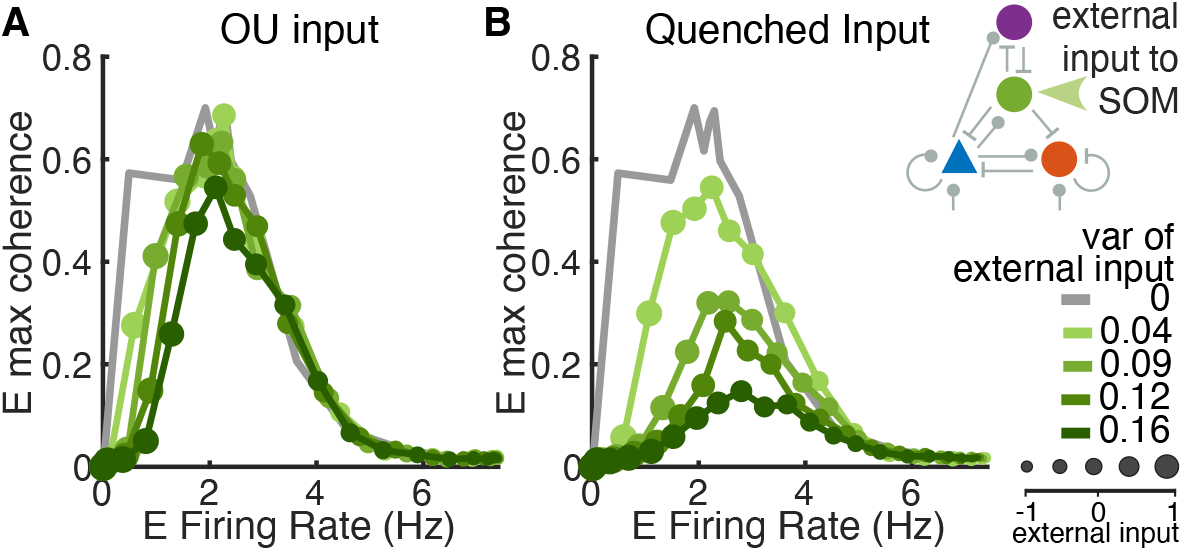
Comparison of two types of external input to SOM neurons. (A) Input to each neuron is modeled as an independent Ornstein–Uhlenbeck (OU-) process with the same mean and variance. (B) Input to each neuron is static in time but the strength is sampled independently from a normal distribution with a specified variance. The case of static input (variance equal to 0) is plotted in grey for comparison in each case (same as the grey curves in Fig. 6).

Interestingly, we find that quenched input to SOM neurons has a much larger impact on network synchrony than OU input with the same variance (Fig. 8). With quenched input, we observe a substantial decrease in E coherence across most firing rates (Fig. 8B). Across all cases of external quenched or OU inputs, the average population rates evolve similarly to the default network with homogeneous static input as input strength is varied (fig. S16, S17). Firing rates change more gradually with increasing variance in the input, especially in the case of quenched input (fig. S17). One difference across input types is that SOM is able to suppress E activity at lower values of quenched input than it can for static input. Overall, these results show that the transition to a synchronized network state resulting from enhancing the activation of SOM neurons is robust against time-varying noisy input that is of similar mean strength across the network, whereas a noise signal that has a spatially-varying strength is more effective at reducing the network synchrony level.

The quenched input in our model is similar to spike threshold heterogeneity, which has been studied in several previous works (*52–55*). Heterogeneity in spike threshold can linearize the input-output response function of neuron populations, increase the excitability of neurons at low input levels, and improve network’s capacity to encode signals (*52–55*). Similar to our results, it has been shown that the spike threshold heterogeneity in the inhibitory neurons reduces oscillations (*52–54*). The impacts of quenched input and time-varying noise were not previously compared, however. Our results demonstrate that quenched input is much more effective at reducing synchrony than time-varying noise.

## Discussion

In this study, using a spatially-structured spiking model of a canonical neural circuit comprising E, SOM, PV and VIP neurons, we demonstrate that SOM neurons are critical for synchronizing neural population activity. As external drive is varied to any target population, the firing rate of SOM neurons is highly predictive of the coherence level that emerges in the E population (Fig. 4C). Without SOM neurons, network synchrony varies much more gradually with the level of input applied to the E-PV subcircuit (fig. S7). The spatial structure of the network is necessary for the gradual transition from asynchrony to strong synchrony via a weak synchrony state, because it allows for the richer spatiotemporal dynamics associated with this transitional regime, consistent with our past work ( (*38*); fig. S14). In addition, we find that when SOM→PV inhibition is strong, the smooth transition through the weak synchrony state is disrupted and the network becomes highly sensitive to input changes (Fig. 5). We further show that the dynamic interaction between E and SOM neurons is a necessary factor in the emergence of SOM-induced network synchrony, as a network in which E and SOM neurons are dynamically uncoupled remains in the asynchronous state even when SOM firing rate is high (Fig. 6).

Our model reproduces previous experimental findings where optogenetic inactivation of SOM neurons led to a reduction in the oscillatory power of the LFP around 30 Hz while inactivation of PV neurons did the opposite (Fig. 3; (*18*,*24*)). Consistent with these experiments (*18*,*24*), SOM neurons contribute to oscillations of lower frequency (15-20 Hz) and PV neurons contribute to oscillations of higher frequency (∼35 Hz) in our model. We identified that the firing rate of SOM neurons is tightly correlated with the overall network synchrony level (Fig. 4C), which is also consistent with the previous experimental finding that the average activity of SOM neurons co-varies linearly with the gamma power of LFP (20-40 Hz) across multiple visual stimulus conditions (Figure S2 in (*17*)). We find that the firing rate and coherence of E neurons can vary in opposite directions through the weak synchrony regime in networks with three interneuron subtypes (Fig. 4A). In contrast, rates and coherence are tethered to vary in the same direction in the E-PV subcircuit (fig. S7). SOM neurons are responsible for the opposite relationship between rate and coherence of E neurons; when SOM neurons are more active, they suppress E neurons and increase network synchrony, and when SOM neurons are suppressed, E neurons firing rate increases and network synchrony is reduced. The opposite directionality of changes in E firing rates versus network synchrony has been observed with changes in spatial attention (*56*) and arousal state (*15, 57*). The simultaneous increase in firing rate and decrease in synchrony can presumably enhance the signal-to-noise ratio of neural representations of stimuli (*58*). Our results suggest that incorporating multiple interneuron subtypes supports the robust emergence of this enhanced coding state.

Our model predicts that a stronger or comparable magnitude of inhibition from SOM to E neurons compared to that from SOM to PV neurons is important for maintaining the weak synchrony regime (Fig. 5, S10). When SOM to PV inhibition is much larger, the network shows abrupt transitions from the asynchronous to the strongly synchronous regime. This sensitivity arises because the positive feedback in the SOM→PV→E→SOM disinhibitory loop can lead to instability. Our result is consistent with a previous model which suggests that SOM inhibition to PV neurons can result in a loss of stability (*59*). The presence of stronger SOM inhibition onto E compared to PV neurons is in agreement with anatomical findings in cortex (*40, 41, 60*). On the other hand, recent experimental work suggests that activating SOM neurons enhances the reliability of E neuron responses to natural movie stimuli by suppressing PV neurons (*29*). The discrepancy between our model and this work could be due to the different temporal patterns of stimulation across the two. In our model, we only consider sustained application of external input, to model slow processes like the variation of brain state, while in these experiments (*29*), pulse stimulation was used. Further analysis is needed to investigate the dynamic responses of our model to brief, cell-type specific stimulation.

Our results also reveal an advantage of targeting VIP neurons to modulate a network’s dynamical state. That is, targeting VIP neurons flexibly transitions the network between asynchronous and weakly synchronous regimes without pushing the network to pathologically strong oscillations (Fig. 3). Anatomically, VIP neurons reside mostly in superficial layers in cortex and receive mostly long-range projections from other brain regions (*9*). Therefore, they are hypothesized to be the main locus of feedback connections and neuromodulator release. VIP neurons also have been shown to respond strongly to locomotion signals (*12*), novel stimuli and unexpected events (*61, 62*). Nevertheless, VIP neurons mainly act through SOM neurons to regulate the E-PV subcircuit. Therefore, it is the activity of SOM neurons that is mostly reflective of network state in our model.

Spatial networks of multiple neuron types have been studied in several modeling works. Some of the previous works focus on analyzing the role of SOM neurons in mediating surround suppression and processing of different visual stimulus patterns (*31, 63–65*). For example, SOM-mediated lateral inhibition is shown to contribute to network responses to edges and motions of visual inputs, and VIP neurons can regulate a network’s processing mode to respond to different features of stimuli (*63, 64*). In a highly detailed microcircuit model of multiple cortical layers of the mouse visual cortex, different tuning-dependent connectivity rules are analyzed to match the distributions of orientation and direction selectivity between models and data (*66*). Most of these works focus on firing rate modulations and the asynchronous dynamical regime. In contrast to these past works, our work focuses on the modulations of rates and network synchrony by cell-type specific top-down input, and only considers spatially uniform feedforward inputs from visual stimuli.

Several recent models also analyze the oscillatory activity in networks of spiking neurons or rate units (*37, 65, 67, 68*). Our work is in agreement with previous results, which suggest that the SOM neurons contribute to oscillations in the low frequency (beta) range and PV neurons contribute to oscillations in the high frequency (gamma) range, though the values of peak frequencies differ across studies. The differences in peak frequencies can be due to different choices of synaptic time scales. When matching the synaptic decay time constants in our model and those used in a previous model (*65*), our network exhibits the same peak frequencies as the earlier model in both gamma and beta ranges. Consistent with previous models (*37, 65*), we also find that the distinct oscillation frequencies associated with PV and SOM neurons, respectively, are largely shaped by their distinct connectivity patterns in that they still exist even when the synaptic time constants of PV and SOM neurons are the same (Fig. 7B-D). The unique contribution in our work is that we systematically analyze the impacts on oscillations and rates from cell-type specific modulatory inputs and reveal common modulation patterns from varying input to E or PV neurons or input to SOM or VIP neurons. We identify the activity of SOM neurons as the main indicator of network synchrony in all modulation cases irrespective of the target of modulatory input, which is not demonstrated in previous works. Our subsequent analysis on the dependence of SOM connectivity provides a comprehensive analysis on SOM-induced network synchrony.

Even though our model considers realistic spatial connectivity patterns and most parameters are constrained by anatomical and physiological data from mouse visual cortex, our model ignores many biological details that can be important for the functions of interneuron subtypes. First, an important distinction between PV and SOM neurons is that PV neurons mainly target the soma of pyramidal neurons while SOM neurons mainly target the dendrites. The dendrite-targeting SOM neurons allow for selectively gating of branch-specific inputs (*69*). Another model also suggests that inputs to VIP neurons can redistribute somato-dendritic inhibition of pyramidal cells and thus control the integration and cancelation of top-down signals that target apical dendrites (*70*). The mutual inhibition between VIP and SOM neurons amplify their responses to small mismatches in their inputs, which can produce prediction errors (*70, 71*). Second, our model does not consider slow physiological mechanisms, such as short-term plasticity, spike frequency adaption and slow synaptic receptors. As a consequence, our model lacks activity fluctuations in low frequency range (*<*10 Hz). Experimental findings suggest that arousal state often has opposing impacts on the low-versus high-frequency oscillatory power of LFP; the high arousal state is associated with reduced power in the low-frequency band and increased power in the high-frequency band (*15, 57, 72*). Additionally, past work has shown that activation of SOM neurons reduces low-frequency power (*<*10 Hz) of LFP in addition to increasing high-frequency power (*10*). Future work is needed to extend the current model to account for the impacts of brain state on population activity across frequencies. Lastly, our network models for the superficial layer of mouse visual cortex, and does not consider the interactions with other cortical layers. Past modeling work suggests that interlaminal connectivity patterns contribute to oscillations of different frequencies (*37, 73, 74*).

The brain features a vast diversity of neuronal types, each of which has unique connectivity patterns, expression of neuromodulator receptors and electrophysiological properties. Different cell types coordinate their activity to regulate neural population dynamics for flexible computations. Our model provides insights and predictions about the different functions that each primary interneuron subtype may serve in modulating the dynamical state of cortex, highlighting the importance of E-SOM interactions and the relative strengths of SOM inputs to E versus PV neurons. Altogether, our results emphasize a unique role of SOM neurons in controlling network synchrony.

## Materials and Methods

### Spiking neuron network model

The model network consists of a single recurrent layer and a feedforward input layer (Fig. 1B). The feedforward layer (population X) is composed of 2,500 (*N*_*X*_) excitatory neurons modeled as independent Poisson processes with a uniform rate of 10 Hz. The recurrent layer contains 50,000 neurons (*N*) divided into four cell population types, *N*_*e*_ = 40, 000 E, *N*_*p*_ = 4, 000 PV, *N*_*s*_ = 4, 000 SOM, and *N*_*ν*_ = 2, 000 VIP neurons. The population size ratios follow anatomical data from mouse cortex (*45*). The synaptic connection patterns among the four neuron populations are constrained by anatomical and physiological data from mouse visual cortex (Fig. 1A; (*25, 40, 41*)). In particular, we assume there are no reciprocal connections among SOM neurons or among VIP neurons; VIP neurons only inhibit SOM neurons; and only E and PV neurons receive input from the feedforward layer (Table 1). Most of model parameters are similar to those in our previous work (*38*) except for some changes to incorporate different interneuron subtypes.

**Table 1.**
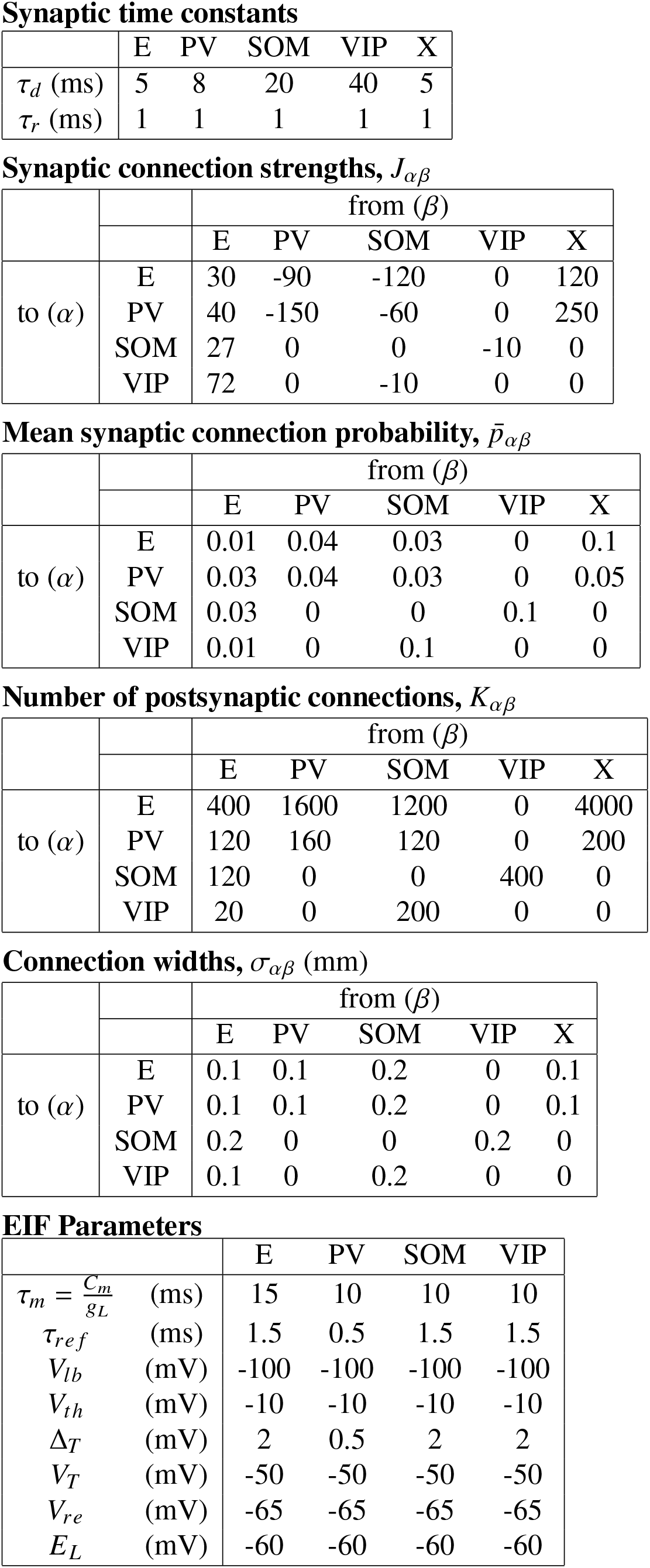
Simulation Parameter Values. The symbol X denotes the feedforward connections.

Synaptic time constants
|  | E | PV | SOM | VIP | X |
| --- | --- | --- | --- | --- | --- |
| $\tau_d$ (ms) | 5 | 8 | 20 | 40 | 5 |
| $\tau_r$ (ms) | 1 | 1 | 1 | 1 | 1 |

Synaptic connection strengths, $J_{\alpha\beta}$
| | | from ($\beta$) | | | | |
| --- | --- | --- | --- | --- | --- | --- |
|  |  | E | PV | SOM | VIP | X |
| to ($\alpha$) | E | 30 | -90 | -120 | 0 | 120 |
|  | PV | 40 | -150 | -60 | 0 | 250 |
|  | SOM | 27 | 0 | 0 | -10 | 0 |
|  | VIP | 72 | 0 | -10 | 0 | 0 |

Mean synaptic connection probability, $\bar{p}_{\alpha\beta}$
| | | from ($\beta$) | | | | |
| --- | --- | --- | --- | --- | --- | --- |
|  |  | E | PV | SOM | VIP | X |
| to ($\alpha$) | E | 0.01 | 0.04 | 0.03 | 0 | 0.1 |
|  | PV | 0.03 | 0.04 | 0.03 | 0 | 0.05 |
|  | SOM | 0.03 | 0 | 0 | 0.1 | 0 |
|  | VIP | 0.01 | 0 | 0.1 | 0 | 0 |

Number of postsynaptic connections, $K_{\alpha\beta}$
| | | from ($\beta$) | | | | |
| --- | --- | --- | --- | --- | --- | --- |
|  |  | E | PV | SOM | VIP | X |
| to ($\alpha$) | E | 400 | 1600 | 1200 | 0 | 4000 |
|  | PV | 120 | 160 | 120 | 0 | 200 |
|  | SOM | 120 | 0 | 0 | 400 | 0 |
|  | VIP | 20 | 0 | 200 | 0 | 0 |

Connection widths, $\sigma_{\alpha\beta}$ (mm)
| | | from ($\beta$) | | | | |
| --- | --- | --- | --- | --- | --- | --- |
|  |  | E | PV | SOM | VIP | X |
| to ($\alpha$) | E | 0.1 | 0.1 | 0.2 | 0 | 0.1 |
|  | PV | 0.1 | 0.1 | 0.2 | 0 | 0.1 |
|  | SOM | 0.2 | 0 | 0 | 0.2 | 0 |
|  | VIP | 0.1 | 0 | 0.2 | 0 | 0 |

|  |  | E | PV | SOM | VIP |
| --- | --- | --- | --- | --- | --- |
| $\tau_m = \frac{C_m}{g_L}$ | (ms) | 15 | 10 | 10 | 10 |
| $\tau_{ref}$ | (ms) | 1.5 | 0.5 | 1.5 | 1.5 |
| $V_{lb}$ | (mV) | -100 | -100 | -100 | -100 |
| $V_{th}$ | (mV) | -10 | -10 | -10 | -10 |
| $\Delta_T$ | (mV) | 2 | 0.5 | 2 | 2 |
| $V_T$ | (mV) | -50 | -50 | -50 | -50 |
| $V_{re}$ | (mV) | -65 | -65 | -65 | -65 |
| $E_L$ | (mV) | -60 | -60 | -60 | -60 |

Each neuron in the recurrent layer is modeled as an exponential integrate-and-fire (EIF) neuron with membrane potential defined as:

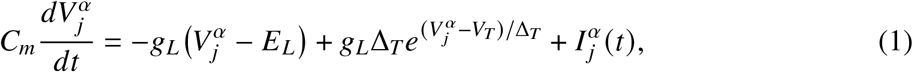

where neuron *j* is a member of the *α* population, *α* ∈ {*e, p, s, ν*}. When 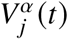 exceeds a threshold *V*_*th’*_ the neuron spikes and the membrane potential is held at *V*_*th’*_ for a refractory period *τ*_*re f*_ and then reset to a lower potential value, *V*_*re*_ (Table 1). All membrane potentials are bounded below by *V*_*lb*_ = −100 mV. The total current to neuron *j* in population *α* is

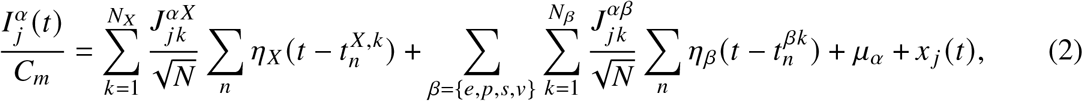

where *n* indexes the spikes fired by the presynaptic neurons, *J*^*αβ*^ is the recurrent synaptic strength from population *β* to population *α* (which may be 0 in some cases), *J*^*αX*^ is the synaptic strength from the feedforward layer to population *α* (Table 1), *µ*_*α*_ is a constant external input current and *x* _*j*_ (*t*) is input noise (Eq. 6). Note that the strength of each synaptic connection is scaled by 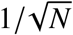. In equation (2), the postsynaptic current terms are defined as:

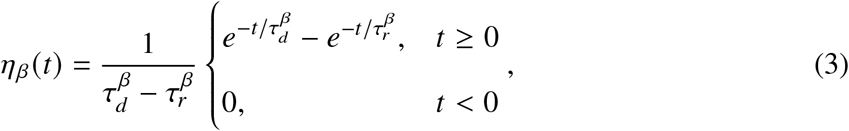

where 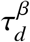 and 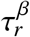 (Table 1) are the synaptic decay and rise time constants for population *β*. The synaptic timescales of inhibitory connections from SOM and VIP neurons are slower than that of connections from PV neurons, which is in turn slower than that of excitatory connections, constrained by physiological data from mouse visual cortex (*44*).

Neurons are uniformly distributed on a unit square, Γ = [0, 1] × [0, 1] (mm^2^). The connection probability between a pair of neurons with coordinates **x** = (*x*_1_, *x*_2_) and **y** = (*y*_1_, *y*_2_), respectively, depends on the populations to which the neurons belong and the distance between the two neurons as

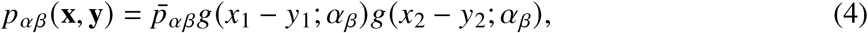

where 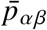 is the mean probability of connections from population *β* to population *α* and *g*(*x*; *σ*) is a wrapped Gaussian distribution:

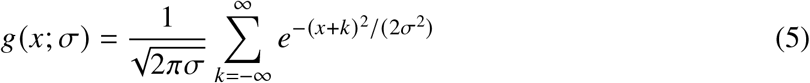

with projection width *σ* (Table 1). Connections to and from the SOM cells have a larger spatial footprint compared to other connections, based on findings from mouse visual and auditory cortex (*25, 43*). A presynaptic neuron is allowed to make more than one synaptic connection to a single postsynaptic neuron. The number of synaptic projections, or out-degree, *K*_*αβ*_, from population *α* to population *β* is fixed for all neurons in population *α*, and indices of postsynaptic neurons are selected randomly according to the connection probability in Eq. 4.

For many of our simulations, the external input, *µ*_*α*_, was varied between -1.0 and 1.0 with step size 0.1. Input noise, *x* _*j*_ (*t*), was modeled as an independent Ornstein-Uhlenbeck (OU) process (Fig. 6, 8):

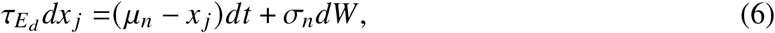

where *W* is a Wiener process, and the time constant of the OU process was chosen to be the same as the decay time constant of the excitatory synaptic current, 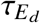 . The mean of *x* _*j*_ (*t*) is *µ*_*n*_ and the variance is 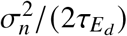. In simulations where we replaced E→SOM connections with an OU process (Fig. 6), we set *µ*_*n*_ = 0.65 and *σ*_*n*_ = 1.1 to match the mean (0.65) and variance (0.12) of the excitatory current from E to SOM neurons in default networks without external input. In simulations with quenched input (Fig. 8B), the constant external input, 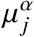, to neuron *j* from population *α* is sampled from a normal distribution with mean *µ*_*α*_ and standard deviation Δ_*µ*_.

The cellular parameters of the EIF model for each cell type and all network parameters are summarized in Table 1. The differential equations (1) and (2) were solved using a forward Euler method with a timestep of 0.05 ms. All simulations were performed on the CNBC Cluster at the Carnegie Mellon University. All simulations were written in a combination of C and MATLAB R2021b (9.11), MathWorks.

### Quantification and Statistical Analysis

#### Spike Count Correlations

Spike counts were computed using a sliding window of 100 ms with a step size of 1 ms. Pearson correlation coefficients were computed for all neuron pairs as a function of distance (Fig. 2iv), except that neurons with rates less than 1 Hz were excluded from correlation calculations. The membrane potential of each neuron was randomly initialized for each simulation, and connectivity matrices were regenerated for each input condition. A total of 5 simulations of 15 seconds each were performed for each input condition. The first 500 ms of each simulation was excluded from the analysis.

#### Coherence

We measured the average pairwise coherence within each cell type population as an indication of network synchrony across simulation conditions. Spike trains were first partitioned into 1 ms time bins and these were collected into 1 second time windows with 0.5 second overlap. Mean firing rate of each sampled neuron was subtracted. Power spectral density, *S*_*i*_, of neuron *i*, and cross spectral density, *S*_*i j*_ , between neuron *i* and neuron *j* , were calculated using the fast Fourier transform and averaged over time windows. The coherence between neuron *i* and neuron *j* at frequency *f* was calculated as

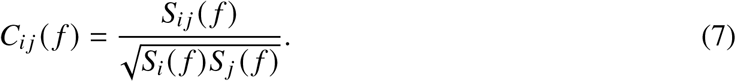

Pairwise coherence was averaged across all sampled neuron pairs within a population. Note that the coherence definition used here is not magnitude-squared, because the magnitude-squared coherence is always positive even when the network is asynchronous. We excluded neurons with rates less than 1 Hz and ensured that 500 neurons were sampled from each population. The first second of each simulation was removed.

#### Activity State Definitions

We identified three network states that were observed for the range of input levels considered, based on mean firing rates and maximum coherence. Specifically, the subcircuit asynchronous (SA) state occurs when the average firing rate of SOM neurons is less than 1 Hz and the maximum coherence of E neurons is less than 0.1. The weakly synchronous (WS) state arises when the maximum coherence of E neurons is between 0.1 and 0.5 and the average firing rate of SOM neurons is larger than 1 Hz. The strongly synchronous (SS) state is when the maximum coherence of E neuron is larger than 0.5.

## Author contributions

Conceptualization: ME, CH, JR

Methodology: ME, CH, JR

Investigation: ME, CH

Formal analysis: ME, CH, JR

Visualization: ME, CH

Data curation: ME, CH

Validation: ME, CH

Software: ME, CH

Project Administration: ME, CH

Funding acquisition: ME, CH

Resources: ME, CH

Supervision: JR, CH

Writing—original draft: ME, CH, JR

Writing—review & editing: ME, CH, JR

## Data and materials availability

All data needed to evaluate the conclusions in the paper are present in the paper and/or the Supplementary Materials. Computer code for all simulations and analysis of the resulting data is available at https://zenodo.org/records/15232542 and https://github.com/hcc11/interneuron/.

## Supplementary Materials

**Figure S1:**
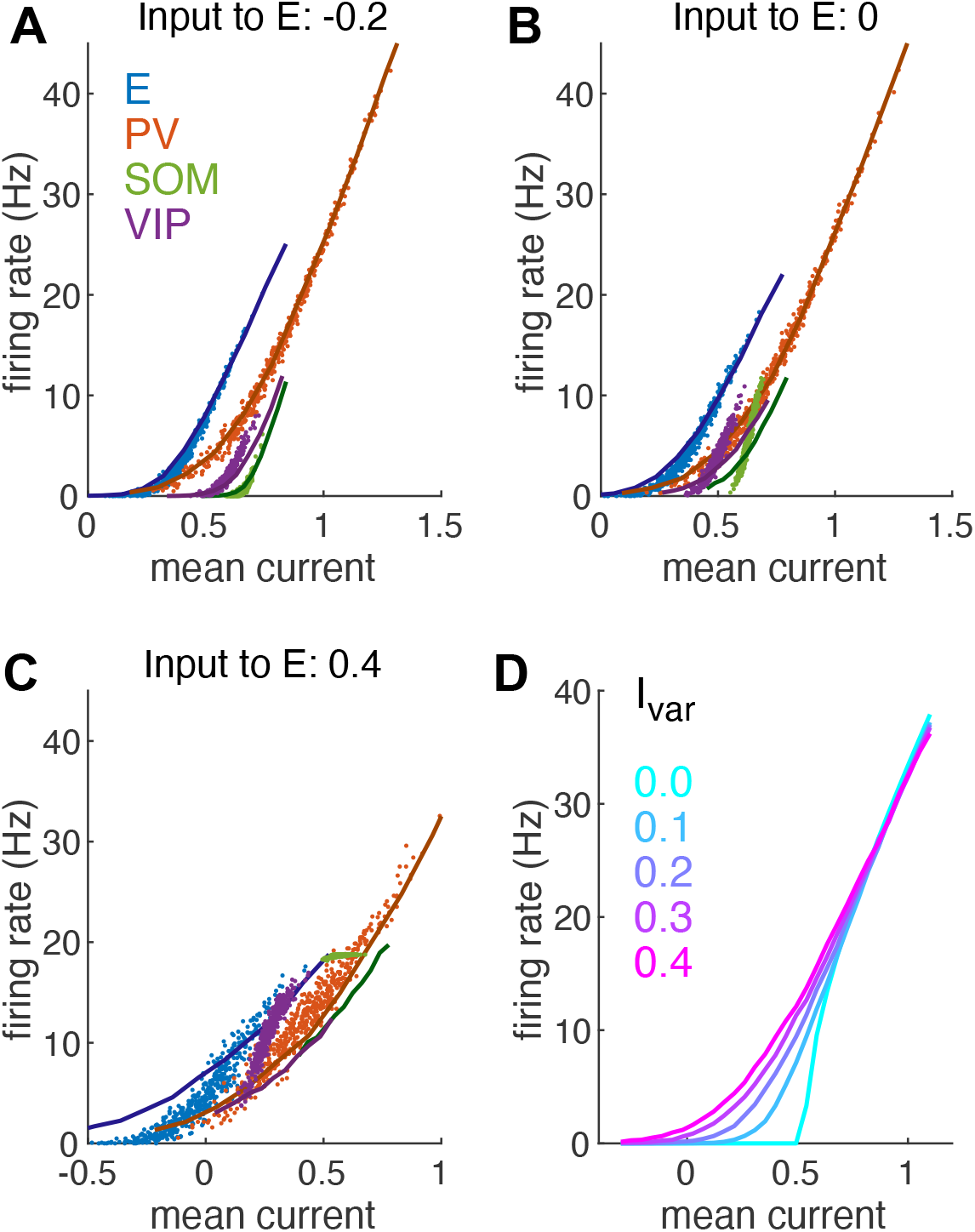
Rate-current transfer functions of each neuron population in the Subcircuit Asynchronous (A), Weakly Synchronous (B) or the Strongly Synchronous (C) state. Each dot represents the mean total current and firing rate of one neuron. There were 500 neurons sampled from each neuron population. Solid curves are the rate-current transfer functions using the population-averaged current variance measured in network simulations of different states and the EIF neuron model parameters for each population. The rate-current transfer functions were calculated by simulating an EIF neuron model (Equation 1 in Methods) with input (*I*(*t*) modeled as an Ornstein–Uhlenbeck process of two time scales: *τ*_*rd*_*I* = (−*I* + *x*)_*d*_*t, τ*_*d*_*dx* = −*xdt* + *JdW*, where *τ*_*r*_ = 1 ms and *τ*_*d*_ = 5 ms were chosen to be the same as those of the excitatory synapses. 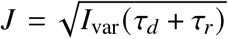 such that the variance of *I* (*t*) is *I*_var_, where *I*_var_ was the population-averaged current variance measured in network simulations. (D) The rate-current transfer function of an EIF neuron with different current variance *I*_var_ (using the E neuron parameters). Note the different x-axis range in C from that in A-B.

**Figure S2:**
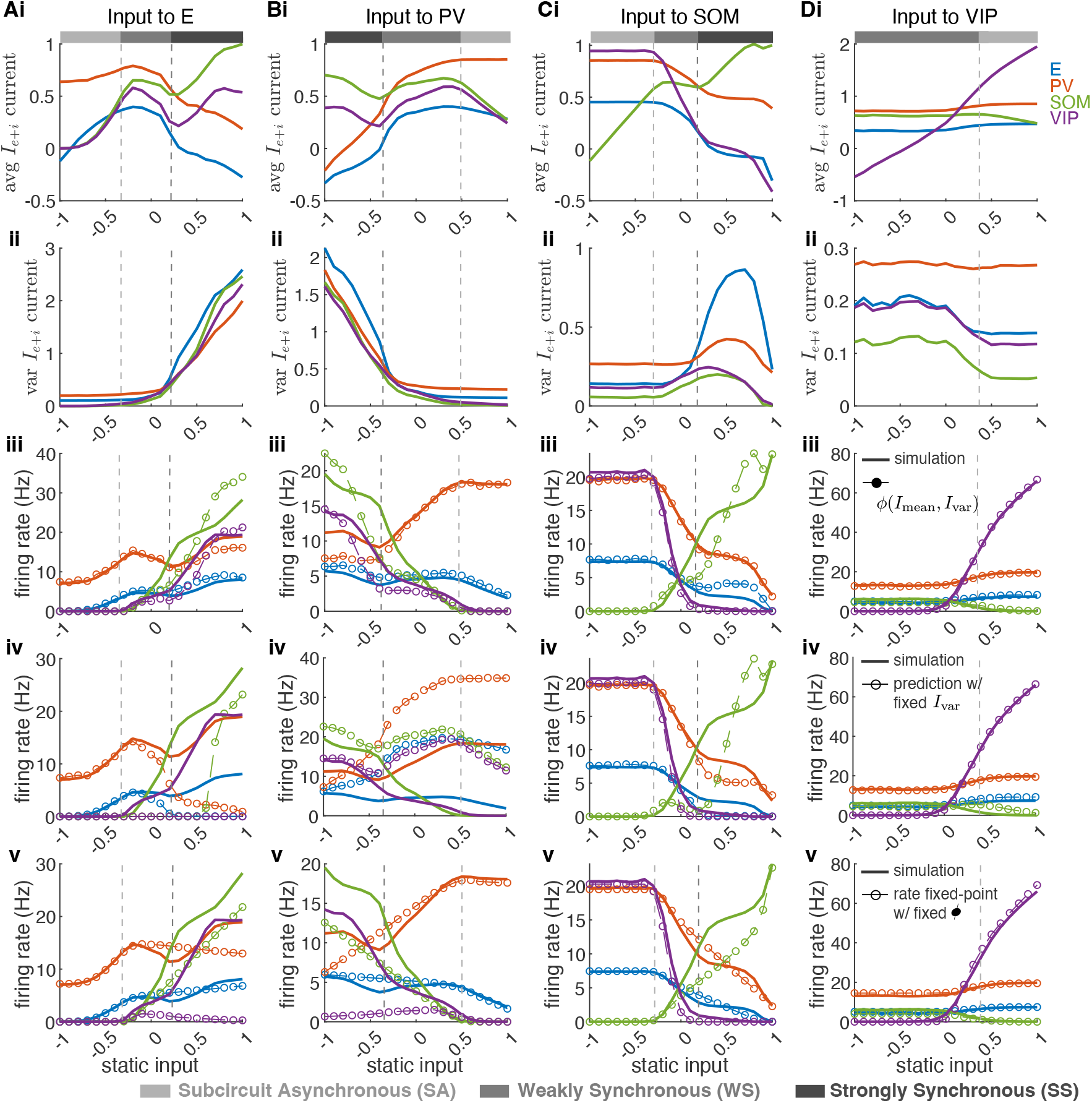
Changes in the mean and variance of synaptic currents and predictions of firing rates as external input is applied to each population. Related to Figure 3. Static input is applied to the E (A), PV (B), SOM (C) or VIP (D) population. Row (i): Average total synaptic current to each population. Row (ii): Population-averaged variance of the total synaptic current to each population. (iii) Dashed curves with circles are predictions of firing rates based on rate-current transfer functions, *ϕ (I*_mean_, *I*_var_) , where *I*_var_ was the population-averaged current variance measured in network simulations (as shown in (ii)). The population firing rate was predicted as the transfer function averaged over the distribution of mean currents (*p (I*_mean_)) from each cell population, i.e. *∫ ϕ* (*I*_mean_, *I*_var_) *p* (*I*_mean_) *dI*_mean_. Solid curves are population firing rates from network simulations, same as in Figure 3Ai-Di. (iv) Dashed with circles: predictions of firing rates, *ϕ* (*I*_mean_, *I*_var_), where *I*_var_ was fixed at the value measured when input = -1 for each population. Solid: same as in iii. (v) Dashed with circles: the solution of the rate equations *r*_*α*_ = *ϕ (∑ w*_*αβ*_*r*_*β*_+ *w*_*αX*_*r*_*X*_+ *µ*_*α*_ ), *α* ∈ {*e, p, s, ν* }, where 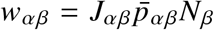 is the effective weight and *r*_*X*_ = 0.01 kHz. The transfer function *ϕ* (*u*) was fixed to be those with *I*_var_ from the asynchronous state (input to E = -0.2 case; Figure S1A). Solid: same as in iii.

**Figure S3:**
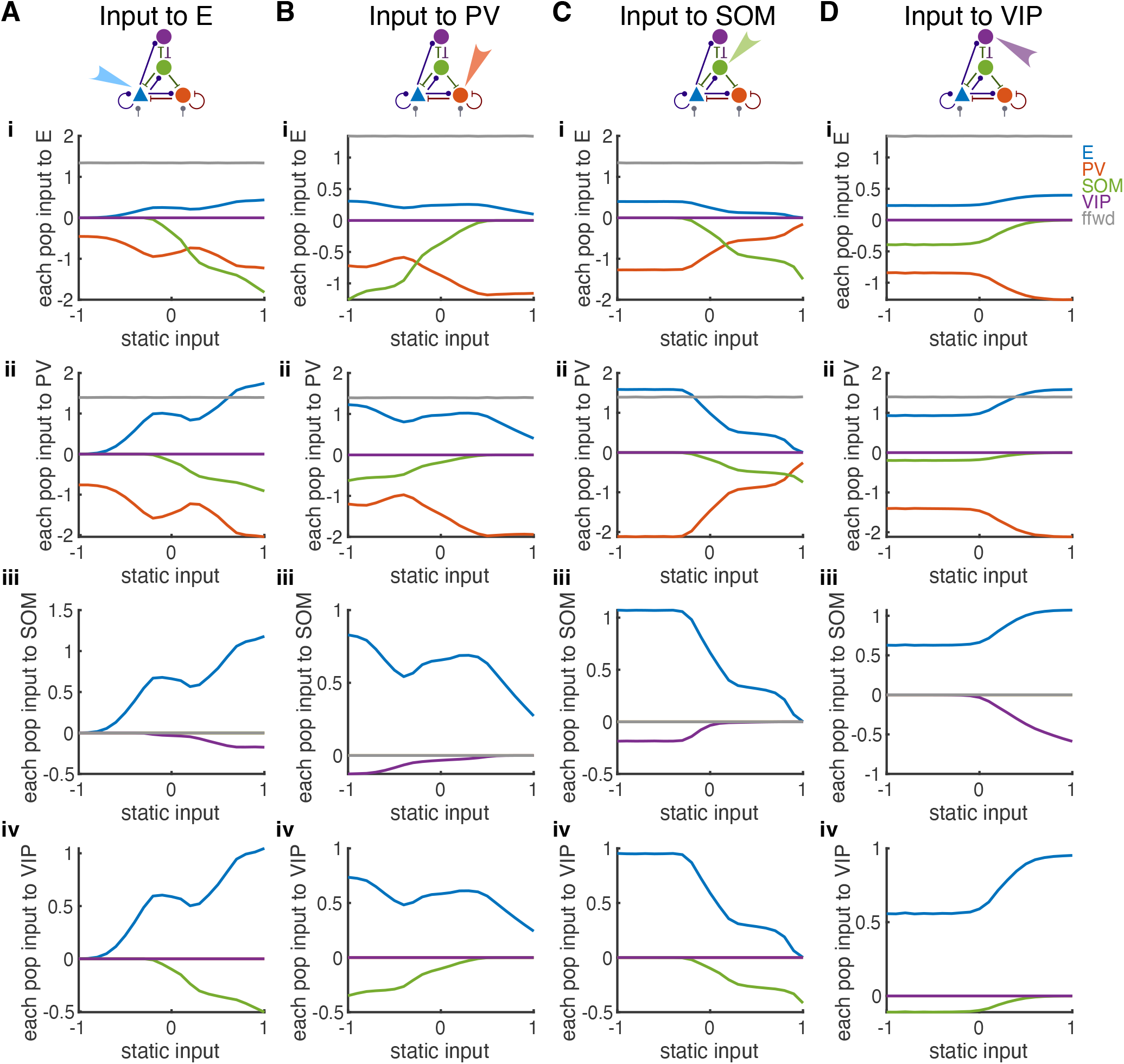
Average synaptic currents from different population sources across modulation states. Related to Figure 3. Static input is applied to the E (A), PV (B), SOM (C) or VIP (D) population. Row (i): average input to the E neurons from E (blue), PV (red), SOM (green), VIP (purple) neurons or neurons in the feedforward layer (grey). The static input was not included. Row (ii): same as (i) for inputs to PV neurons. Row (iii): same as (i) for inputs to SOM neurons. Row (iv): same as (i) for inputs to VIP neurons. The average input from population *β* is 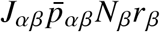, which is proportional to the rate of population *β*. Therefore, inputs from the same population (curves of the same color across rows) have the same shape but with different coefficients.

**Figure S4:**
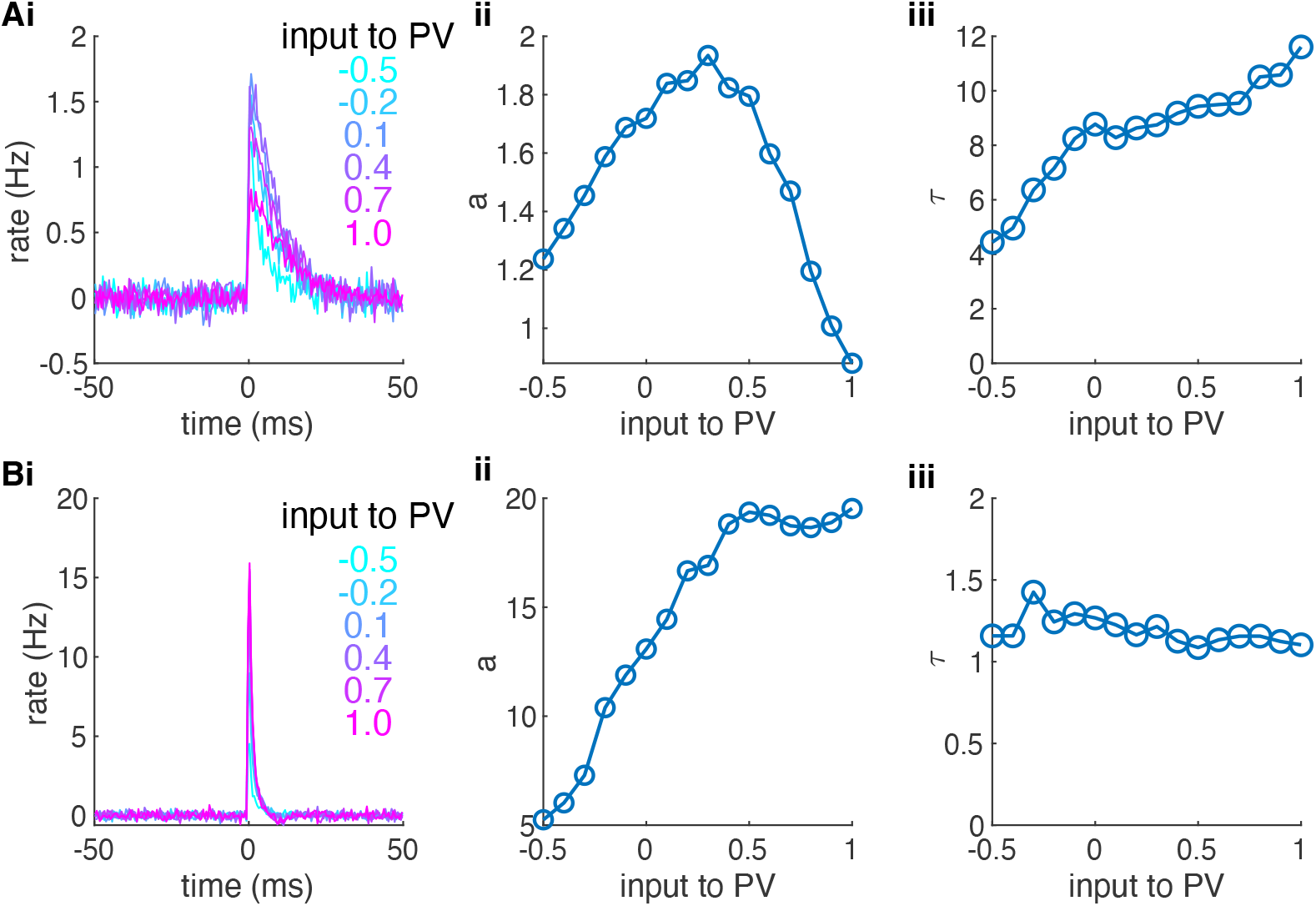
Linear response kernels of the E (A) and the PV (B) neurons with various levels of input applied to the PV neurons. (i): The average firing rate response of individual uncoupled E or PV neurons to an impulse input current at time zero (0.5*δ*(*t*)), which is denoted as linear response kernels (*h (t)* ). The neurons receive colored noise, *I* (*t*) , modeled as an Ornstein–Uhlenbeck (OU) process of two time scales: *τ*_*r*_ *dI* = (−*I* +*x* )*dt, τ*_*d*_*dx* = −*xdt* +*JdW* , same as in Figure S1A-C. *τ*_*r*_ = 1 ms and *τ*_*d*_ = 5 ms were chosen to be the same as those of the excitatory synapses. The mean of the OU noise matched the distribution of mean total current over each neuron population and the variance of the noise matched the population-averag<u>ed current va</u>riance measured in network simulations for each external input case. *J* was set to 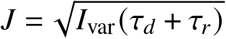 such that the variance of *I* (*t*) was *I*_var_. Each linear response kernel in (i) was fitted with an exponential function, *ae*^−*t*/*τ*^, *t >* 0 . The fitted parameters, *a* and *τ*, are shown in (ii) and (iii), respectively, for different external input values to PV. We find that the response gain (*a*) of PV neurons increases by several folds as input to PV neurons increases (Bii), which means that small fluctuations in the input current to PV result in larger response in PV population rate when PV neurons receive more external inputs. Meanwhile, the response gain of the E neurons first increases and then decreases when the network is about to transition to the asynchronous state (Aii). The reduction in the response gain of the E neurons also contributes to the desynchonization of network dynamics.

**Figure S5:**
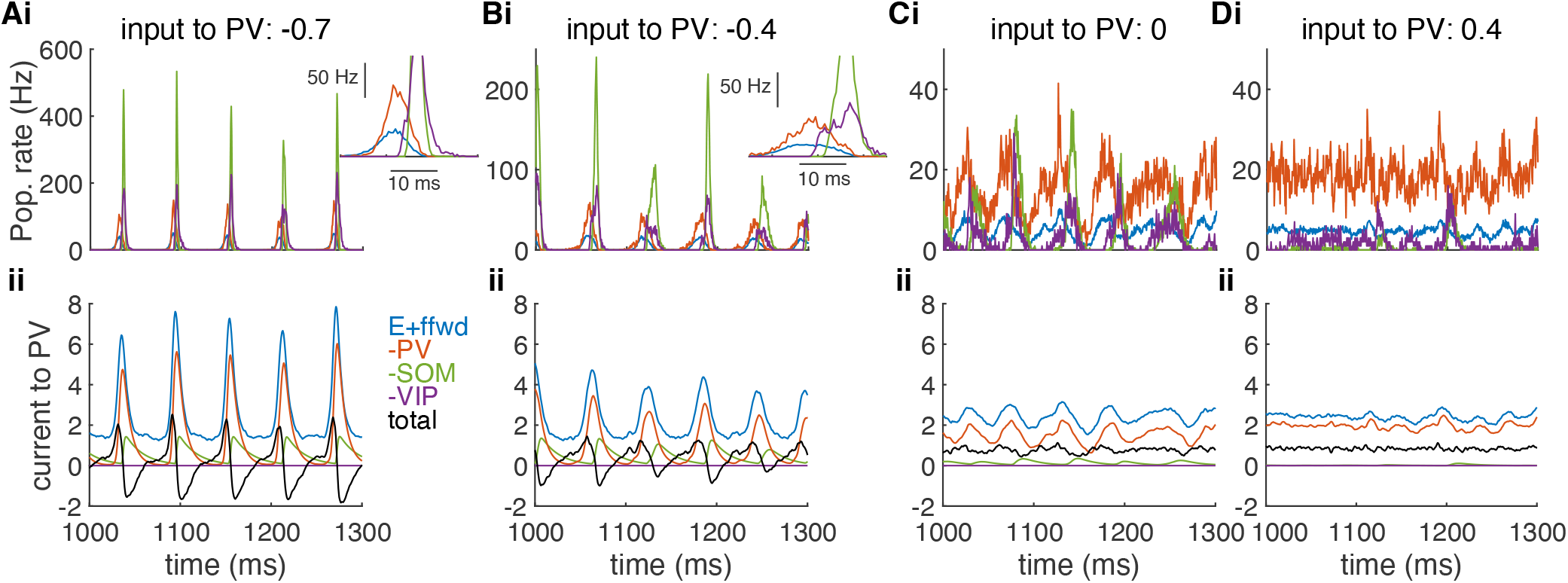
Example traces of firing rates and input currents in different dynamical states. The static inputs applied to PV neurons are -0.7 (A), -0.4 (B), 0 (C) or 0.4 (D). Row (i): Population-average rates of E (blue), PV (red), SOM (green) and VIP (purple) neurons. Note that the y-axis has different ranges in Ai-Di. Insets in Ai and Bi: Zoomed-in view of the population rates during one oscillation cycle. Row (ii): Population-averaged input currents from different sources to PV neurons. Blue: excitatory currents from E neurons and feedforward projecting neurons. Red: absolute value of the inhibitory current from PV neurons. Green: absolute value of the inhibitory current from SOM neurons. Purple: absolute value of the inhibitory current from VIP neurons. Black: total input current. When the input to PV neurons is very negative, the total current to PV neurons remains at a low baseline at the end of an oscillation cycle (Aii, black), which makes the PV neurons lag behind E neurons slightly at the beginning of the next cycle (Ai, inset). A higher input value to PV neurons increases the baseline current level for PV neurons at the end of each cycle, which allows their firing rate to rise in-phase or earlier than the E neurons at the beginning of each cycle (Bi, inset). The earlier firing of PV neurons suppresses the peak of E firing rate and makes the spiking of E neurons more spread out within a cycle. The spread of E spiking reduces the coherence level and reduces the recruitment of SOM neuron activity. Importantly, an increase in PV firing rate also suppresses PV neurons themselves, because of the strong coupling within the PV population. Therefore, PV neurons tend to fire in tandem with E neurons and they do not exert excessive inhibition like the SOM neurons.

**Figure S6:**
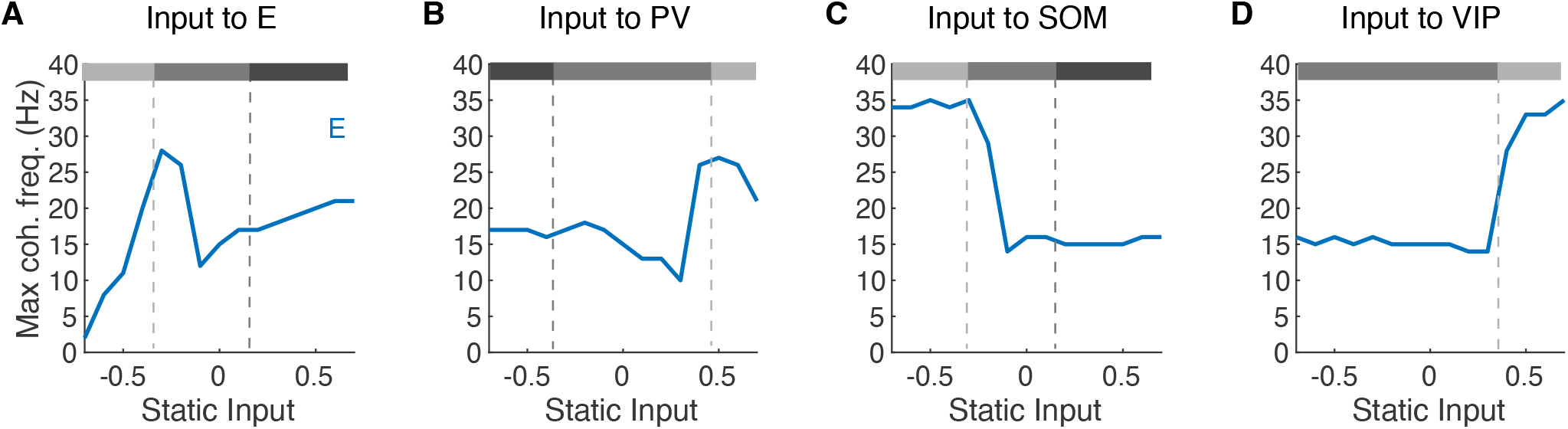
The frequency of maximum coherence of the E neurons across modulation input values. Related to Figure 3. Input is applied to E (A), PV (B), SOM (C) or VIP (D) neurons. Grey-scale bars above each plot represent network activity state at the corresponding input value (Subcircuit Asynchronous: light grey; Weakly Synchronous: moderate grey; Strongly Synchronous: dark grey).

**Figure S7:**
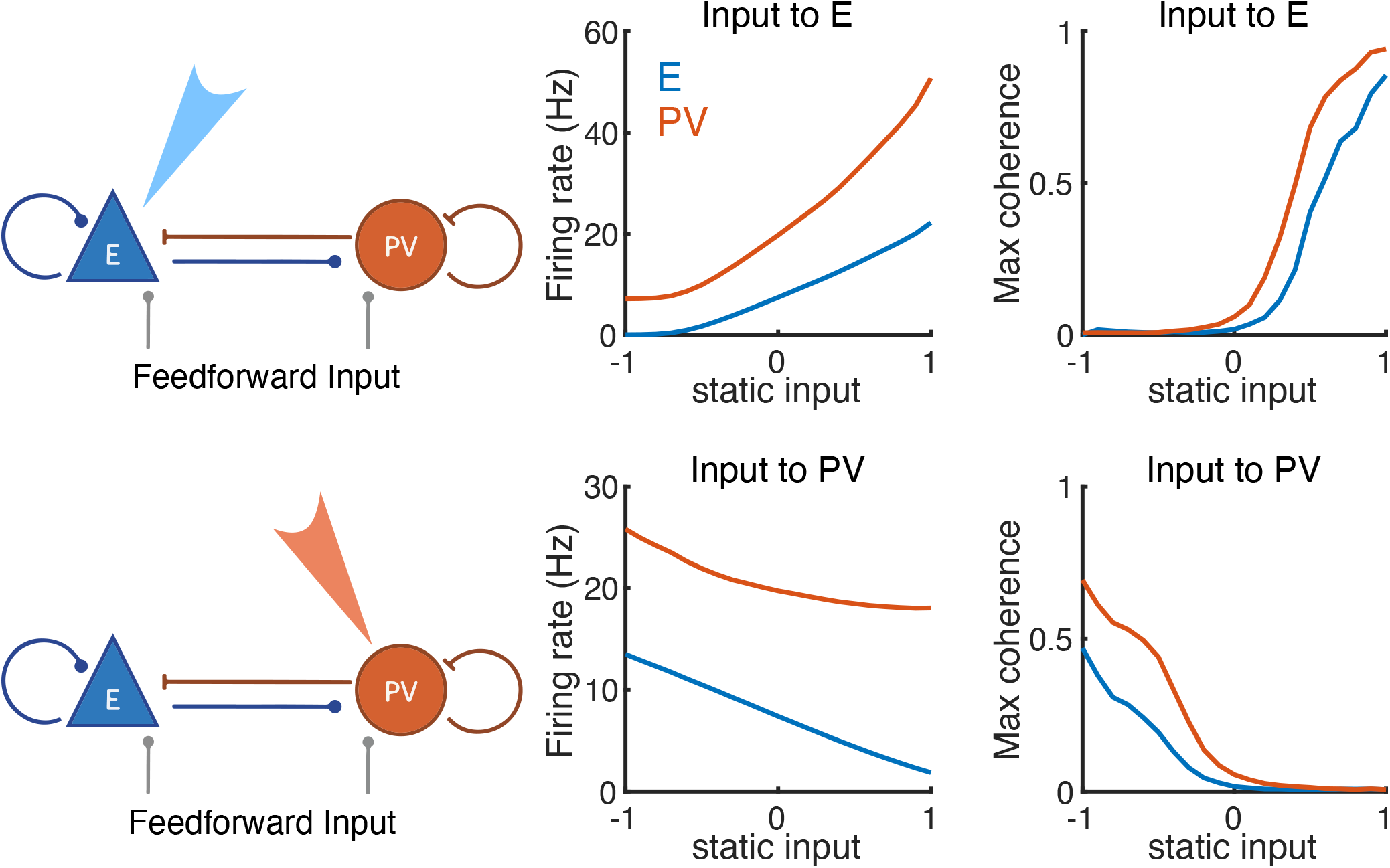
Modulations of firing rates and maximum coherence in the E-PV subcircuit. Related to Figure 4. Static external input was targeted to E (top row) or PV (bottom row) neurons. Firing rates and network synchrony are tethered to change in the same direction in the E-PV subcircuit. That is, stimulating E neurons increases the firing rates and coherence of both E and PV neurons, while stimulating PV neurons decreases firing rates and coherence in both populations. The paradoxical effect where stimulating PV leads to a reduction in PV firing rate suggests that the E-PV subcircuit is in the inhibition-stabilized regime (*75–77*). Network parameters were the same as those Figures 1-4 except that we removed SOM and VIP populations.

**Figure S8:**
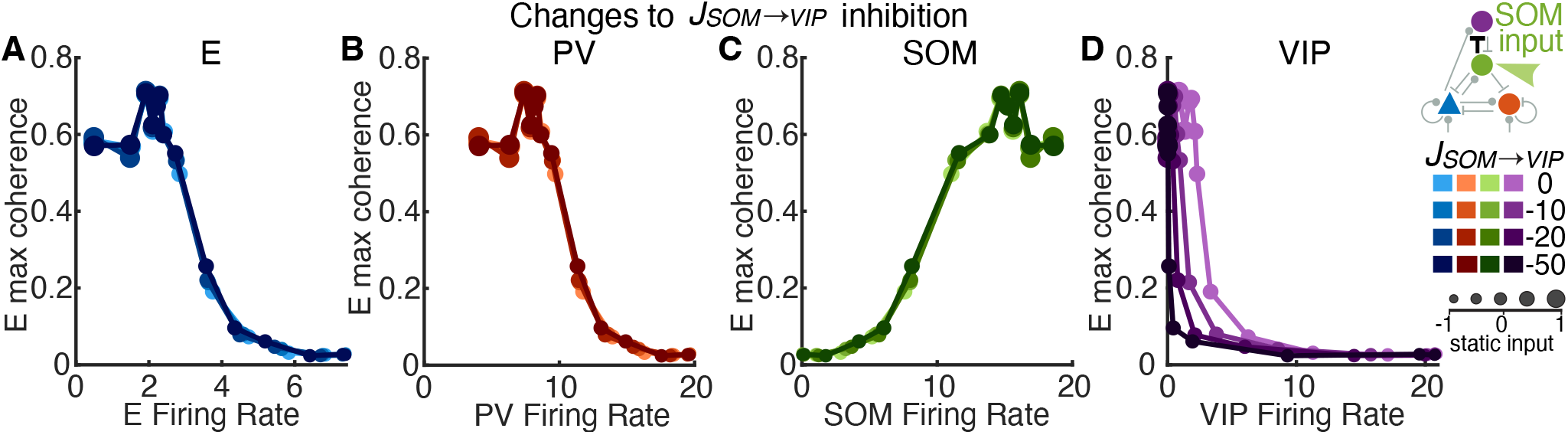
SOM →VIP connection strength has little effect on modulation patterns. Related to Figure 5. Static input was applied to SOM neurons.(A) E, (B) PV, (C) SOM, and (D) VIP population rates compared to E maximum coherence in networks with different SOM →VIP connection strengths, *J*_*SOM* →*VIP*_. Only the relation to VIP firing rate (panel D) is different in networks with different *J*_*SOM* →*VIP*_. In networks with larger *J*_*SOM* →*VIP*_ inhibition, SOM is able to suppress VIP at a lower rate, resulting in the darkened curves shifting leftward (D). Therefore, altering the connection strength of *J*_*SOM* →*VIP*_ exclusively affects the VIP population and does not influence how the rest of the network responds to external input. Note that *J*_*SOM* →*VIP*_ = −10 is the default circuit parameter used in the main text.

**Figure S9:**
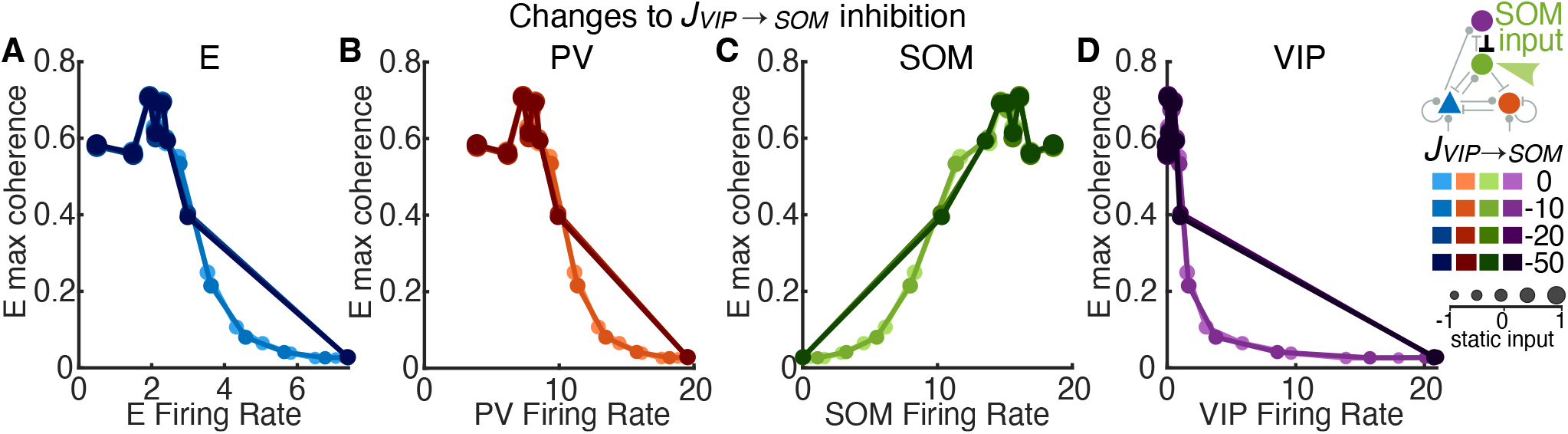
VIP →SOM connection strength has little effect on modulation patterns. Related to Figures 5, 6. Same format as fig. S8. Static input was applied to SOM neurons. Modulation patterns are the same across VIP →SOM connection strengths *J*_*VIP* →*SOM*_ except for in the network with the largest strength (darkest color). Networks with large *J*_*VIP* →*SOM*_ become sensitive to small changes in external input to SOM. The firing rate of SOM neurons switches from zero to above 10 Hz and the rate of VIP neurons switches from about 20 Hz to near zero as input to SOM increases slightly (one step in panels C,D). Therefore, networks with large inhibition from VIP to SOM exhibit the WS state over only a limited parameter range and switch relatively abruptly between the SA and the SS states as input varies. Note that *J*_*VIP* →*SOM*_ = −10 is the default circuit parameter used in the main text.

**Figure S10:**
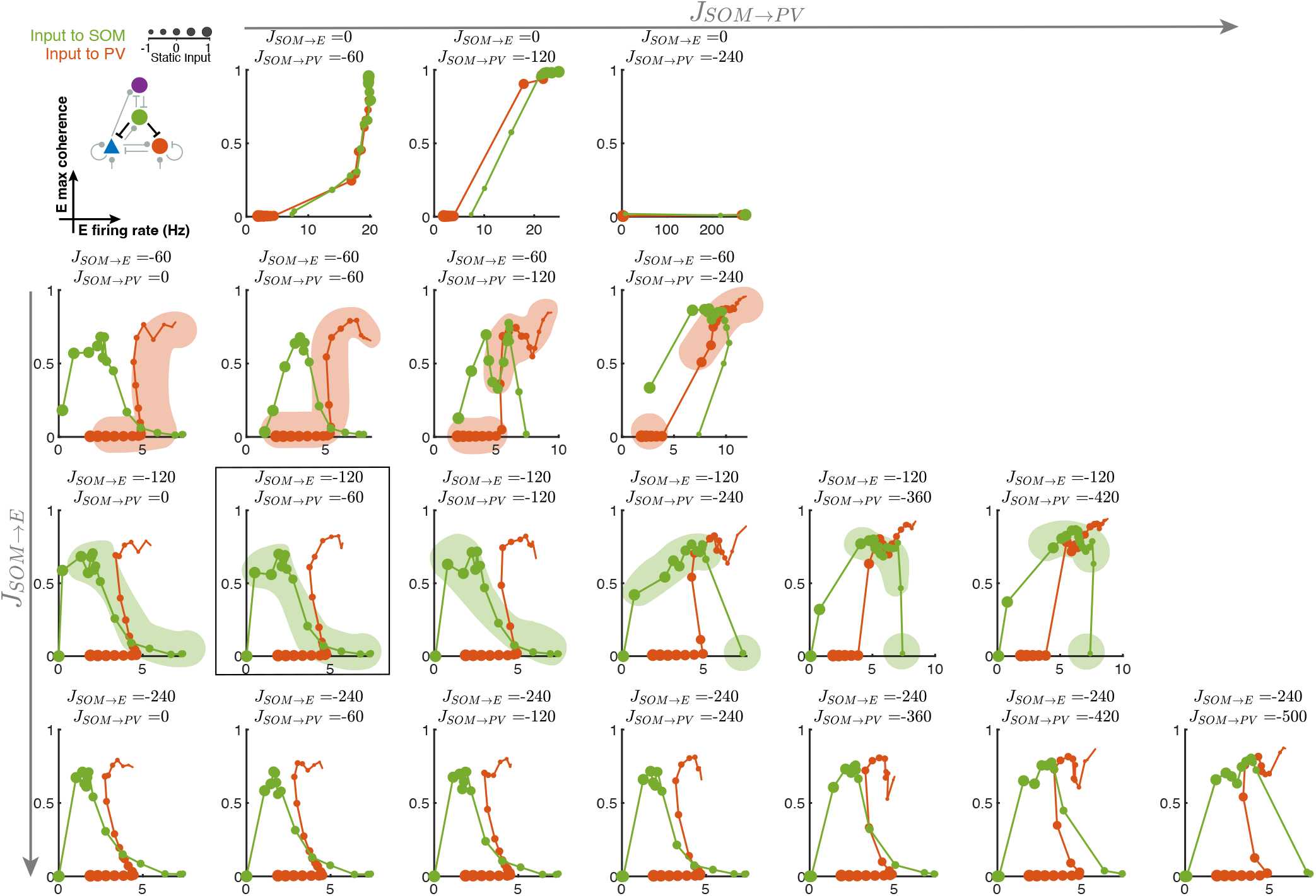
Modulation patterns in networks with different SOM →E and SOM →PV connection strengths. Related to Figure 5. Static input was either applied to PV neurons (red) or SOM neurons (green). Each plot represents the maximum coherence of E neurons versus the average firing rate of E neurons. Marker sizes correspond to increasing static input to the target population. Rows represent (negative) increases in synaptic connection strength of SOM →E. Columns represent (negative) increases in synaptic connection strength of SOM→PV. The boxed plot features the same parameters as the default network. We find that when |*J*_*SOM*→ *E*_ |*>* |*J*_*SOM* →*PV*_ | (lower triangle of the plots), the shapes of the modulation patterns for both input cases remain qualitatively consistent. When |*J*_*SOM* →*PV*_| is much larger than |*J*_*SOM*→ *E*_| , the network exhibits abrupt changes from the SA to the SS state, and the firing rate and coherence of E neurons tend to vary in the same direction over all levels of input to PV. The orange shading in row two and green shading in row three highlight examples of changes in modulation patterns as |*J*_*SOM*→ *PV*_ |increases (across columns in the same row). Discontinuities in the shading are abrupt changes between adjacent dots (i.e., small changes in external input leading to large changes in coherence) along the modulation path.

**Figure S11:**
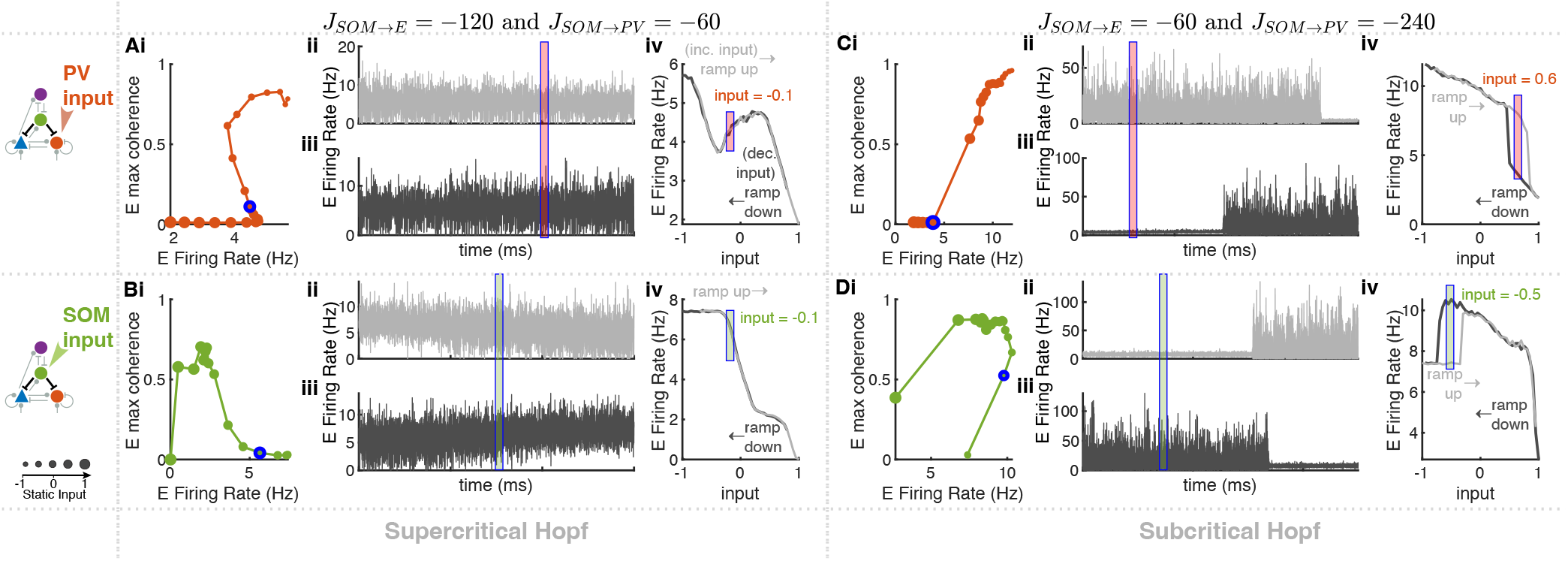
Hysteresis effects in networks with strong SOM →PV inhibition in response to ramping input. Related to Figures 3,4,5. (A,B) The default network (*J*_*SOM* →*E*_ = −120 and *J*_*SOM* →*PV*_ = −60; Figures 3,4) with input applied to PV (A) or SOM (B) neurons. (C,D) Same as A,B for a network with stronger SOM to PV inhibition (*J*_*SOM* →*E*_ = −60, *J*_*SOM* →*PV*_ = −240) with input applied to PV (C) or SOM (D) neurons. Panel (i): Modulation path of E population firing rates versus E maximum coherence with varying static input (same format as Figures 4A, S10). The blue outlined marker in (i) represents the input value that is indicated in panels (ii)-(iv) by colored rectangles. Note that each panel (i) was generated with a sequence of fixed values of static input to the indicated population, and not with ramping input, which changes in time. Panel (ii): The population-averaged firing rate of E neurons as a function of time with slowly increasing input (*ramp up* case). Panel (iii): same as panel (ii) for slowly decreasing input (*ramp down* case). In the ramping input cases, external inputs were increased or decreased by a small incremental change, ± 0.05, every 5 seconds. The 5-second interval allows sufficient time for the network to converge to a stationary state at the given input value. The colored rectangle in panels (ii) and (iii) indicates time intervals of the same input value in both ramping cases, which are aligned in time for comparison. Panel (iv): E population-average firing rates calculated within each 5-second interval of ramping input. The initial 250 ms of each interval was excluded to avoid transient activity. In the default network with |*J*_*SOM*→ *E*_ |*>*| *J*_*SOM* →*PV*_| (A,B), the E firing rate is the same for each fixed input value in both *ramp up* and *ramp down* cases. This suggests that there is no co-existence of multiple network states for any input value and that the transition from the SA to the SS state is likely through a supercritical Hopf bifurcation where the amplitude of oscillation increases gradually after bifurcation. In contrast, in a network with |*J*_*SOM*→ *E*_ |*<* |*J*_*SOM* →*PV*_| (C,D), the same input value results in different dynamic states in the *ramp up* and *ramp down* cases (Cii-Civ,Dii-Div, regions indicated by colored rectangles). This hysteresis effect demonstrates the co-existence of two network solutions, one asynchronous and one synchronous oscillation, over a range of input values. This suggests that oscillations arise via a subcritical Hopf bifurcation, where there is a sudden jump in the amplitude of oscillations after the bifurcation point, in networks with strong SOM→PV inhibition.

**Figure S12:**
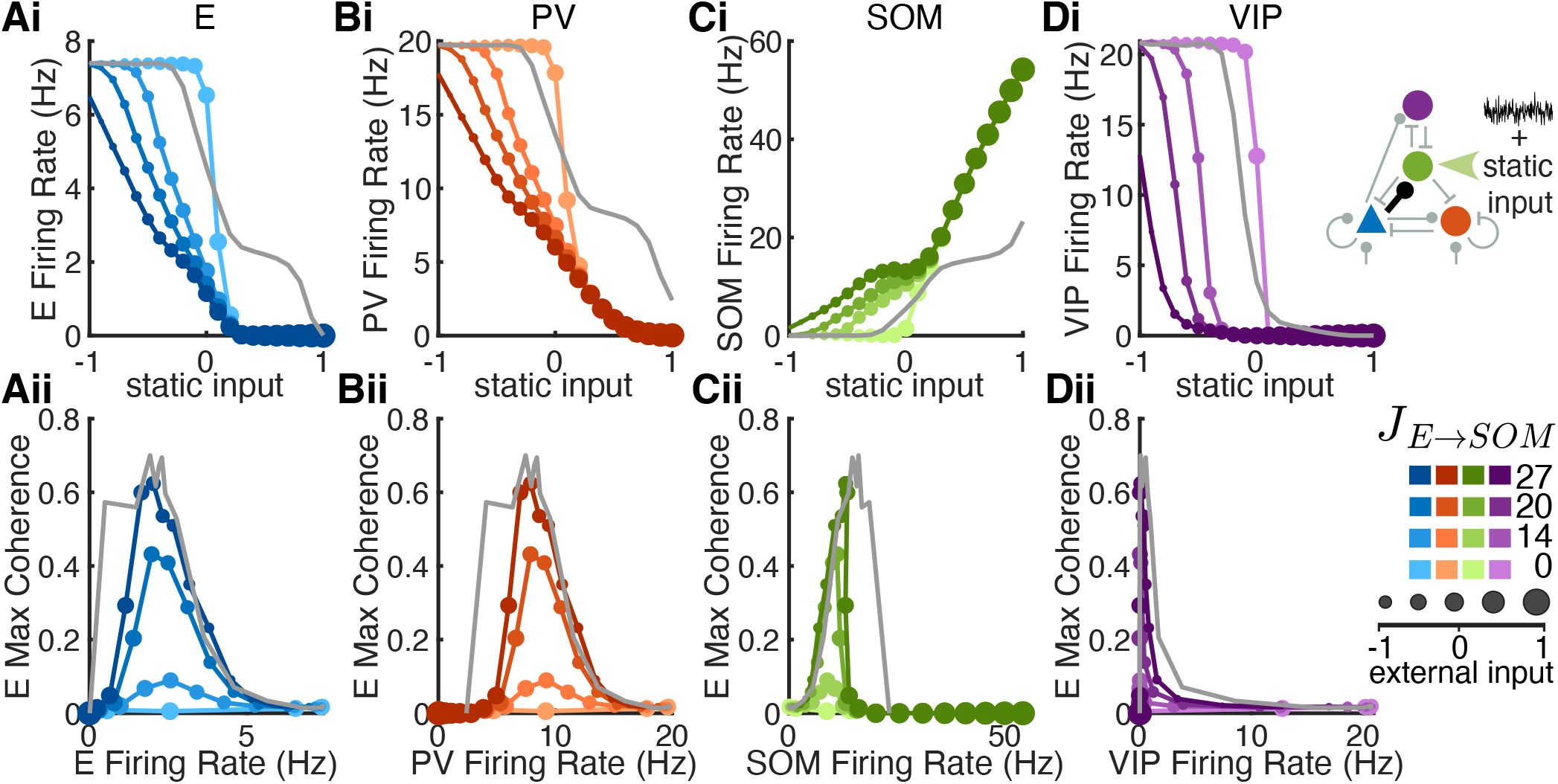
Population rates and coherence in networks with different E →SOM connection strengths. Related to Figure 6. Both static input and colored noise were applied to SOM neurons. The colored noise was constructed as an OU process to match the mean and variance of the recurrent excitatory input that SOM neurons receive in the default network without external input (same noise input as in Figure 6A,C). Static input varied from -1 to 1. Column: (A) E, (B) PV, (C) SOM, (D) VIP population. Row (i): Average firing rates of each cell population with respect to static input value. Row (ii): The maximum coherence of E neurons compared to the population firing rates of each cell type. The grey curves are from the default network (*J*_*E* →*SOM*_ = 27), with SOM neurons receiving static input without the OU noise.

**Figure S13:**
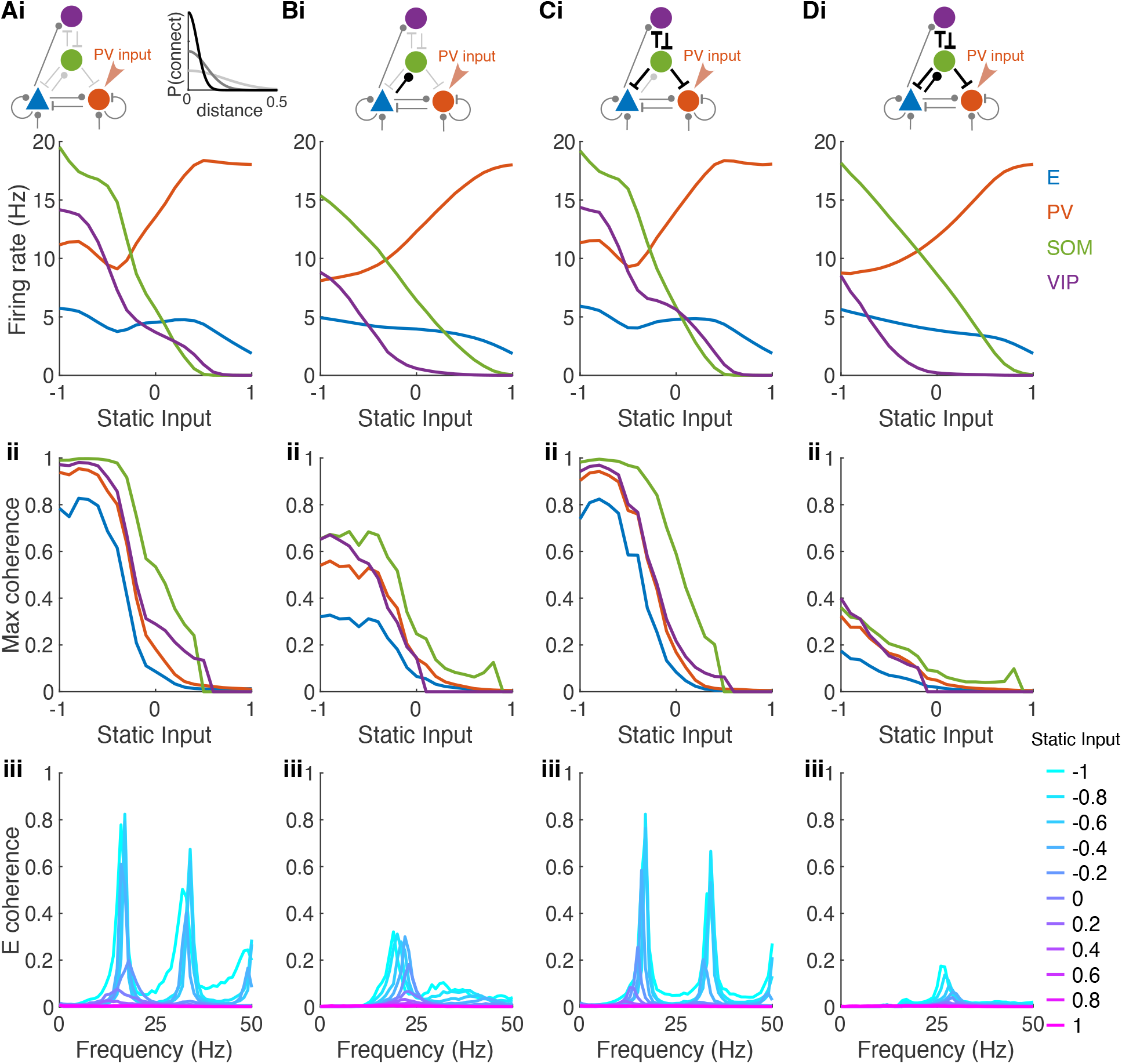
Impacts of the projection widths of SOM connections on network synchrony. Related to Figure 7A. Networks in A-D are the same as those in Figure 7Ai-iv, respectively. Row (i): Average population firing rates of each population as a function of static input value. Row (ii): Maximum coherence in each population as a function of static input value (same as Figure 7A). Row (iii): E population coherence as a function of frequency for several static input values. (A), default network where connections to and from SOM neurons were broader (width = 0.2 mm) compared to other connections (width = 0.1 mm). (B), E →SOM connection width is narrowed to 0.05 mm while keeping all other connection widths the same as those in the default network. (C), SOM connections other than E→ SOM, i.e. VIP→ SOM, SOM→ E, SOM →PV and SOM→ VIP connections, are narrowed to 0.05 mm while keeping all other connection widths the same as those in the default network. (D), all connections from and to SOM neurons are narrowed to 0.05 mm, while keeping all other connection widths the same as those in the default network. In all four networks, static input was applied to the PV neurons and varied from -1 to 1.

**Figure S14:**
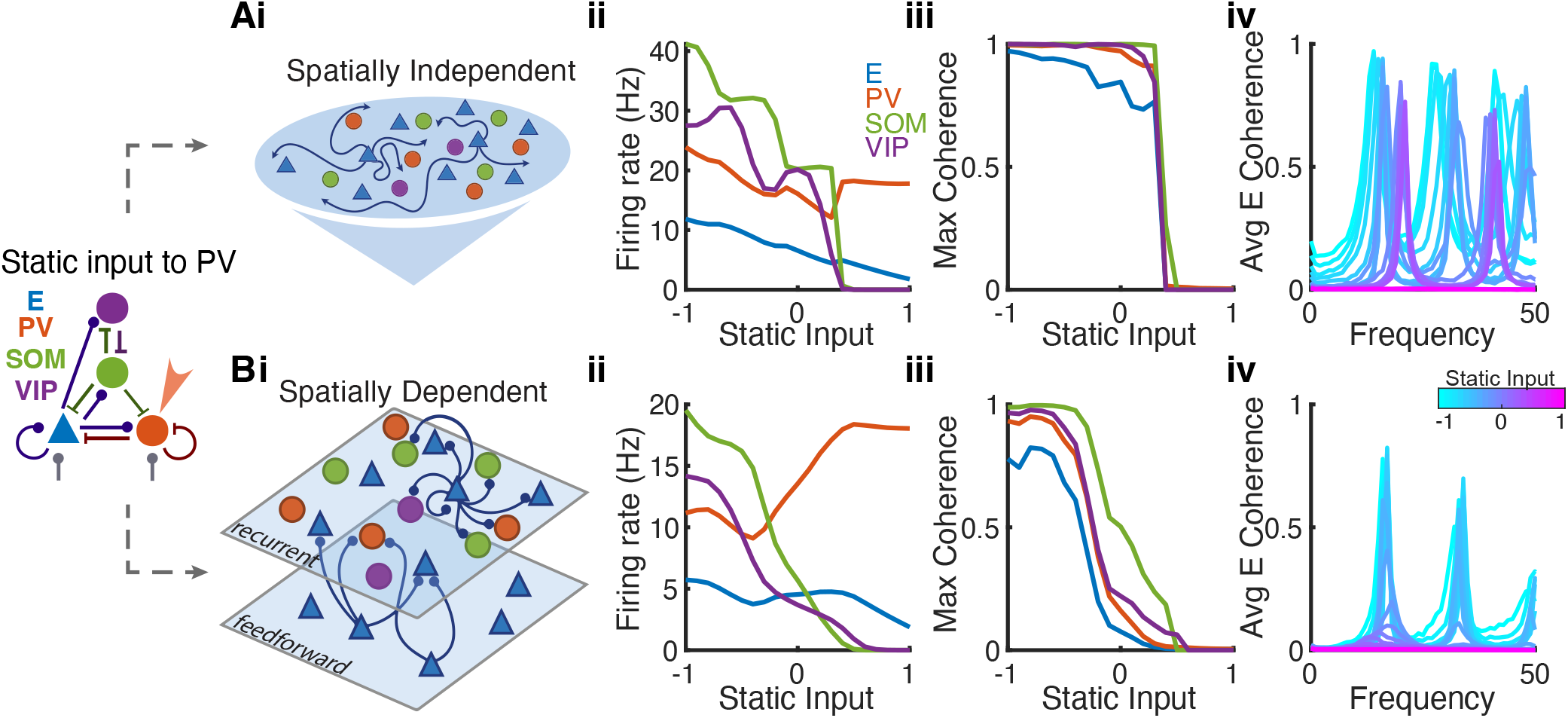
Sharp transitions in networks with no spatial structure. Related to Figures 3,7. Static input was applied to PV neurons. (Ai-iv) Firing rate and coherence in the four neuron populations in networks with no spatial structure, meaning that the connection probability between two neurons does not depend on distance. A sharp transition from the asynchronous to the strongly synchronous state occurs as input is increased. (Bi-iv) The same quantities for networks with spatial structure, as also shown in Figure 3B. The spatially dependent network exhibits a more gradual transition and the existence of weakly synchronous state over a range of input values. The parameters of the networks in A and B are the same except for the connection widths, *σ*_*αβ*_ (Table 1).

**Figure S15:**
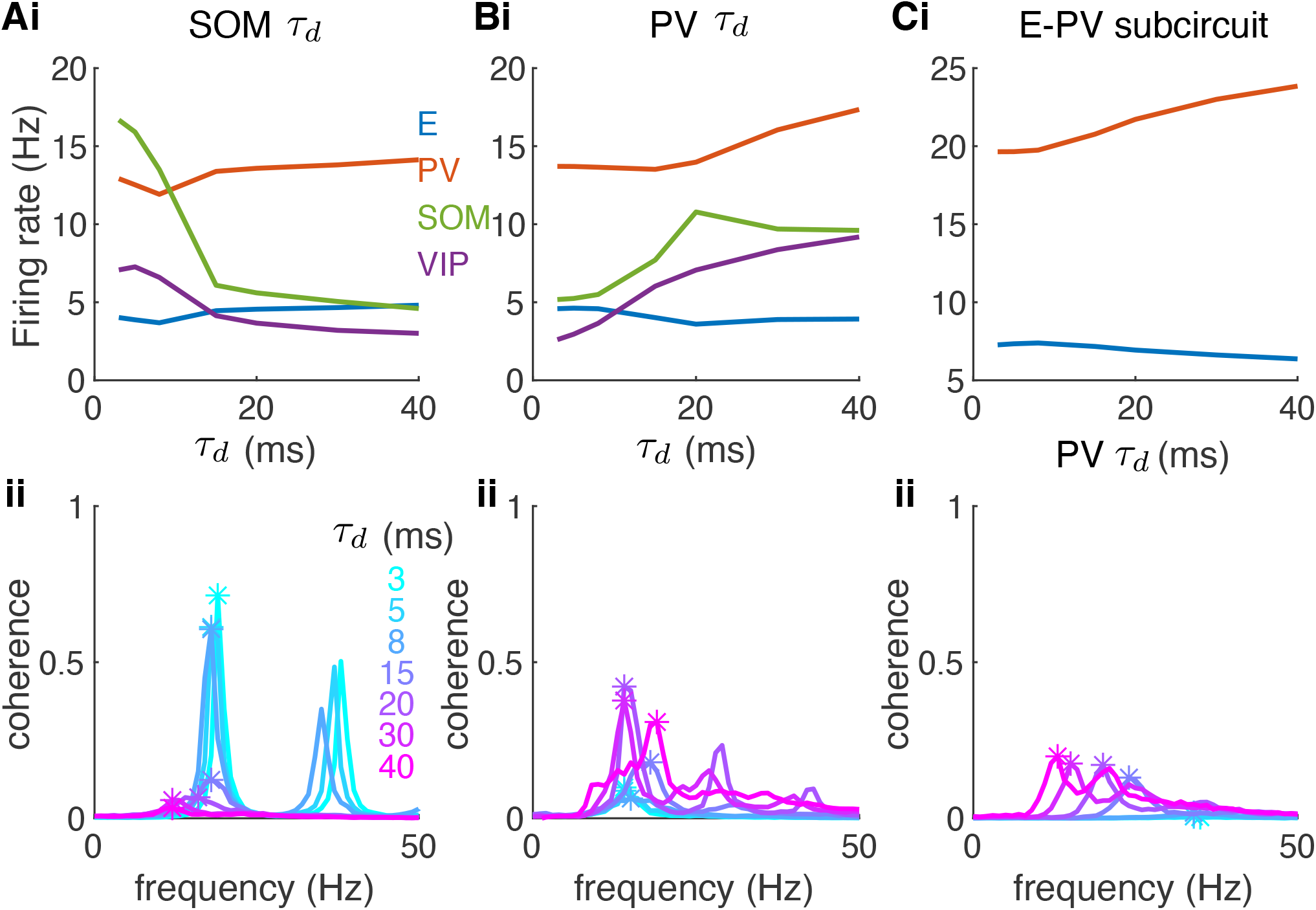
Impacts of the synaptic time scales on rates and network synchrony. Related to Figure 7B-D. (A) Varying the decay time constant (*τ*_*d*_) of SOM synapses. (B) Varying the decay time constant of PV synapses. (C) Same as B in the E-PV subcircuit. Row (i): Average population firing rates of each population as a function of *τ*_*d*_. Row (ii): E population coherence as a function of frequency for networks with different values of *τ*_*d*_. Other parameters were the same as those in Figure 3. The static input was zero. For reference, *τ*_*d*_ is 8 ms and 20 ms for the PV and SOM synapses, respectively, in the default network in Figure 3.

**Figure S16:**
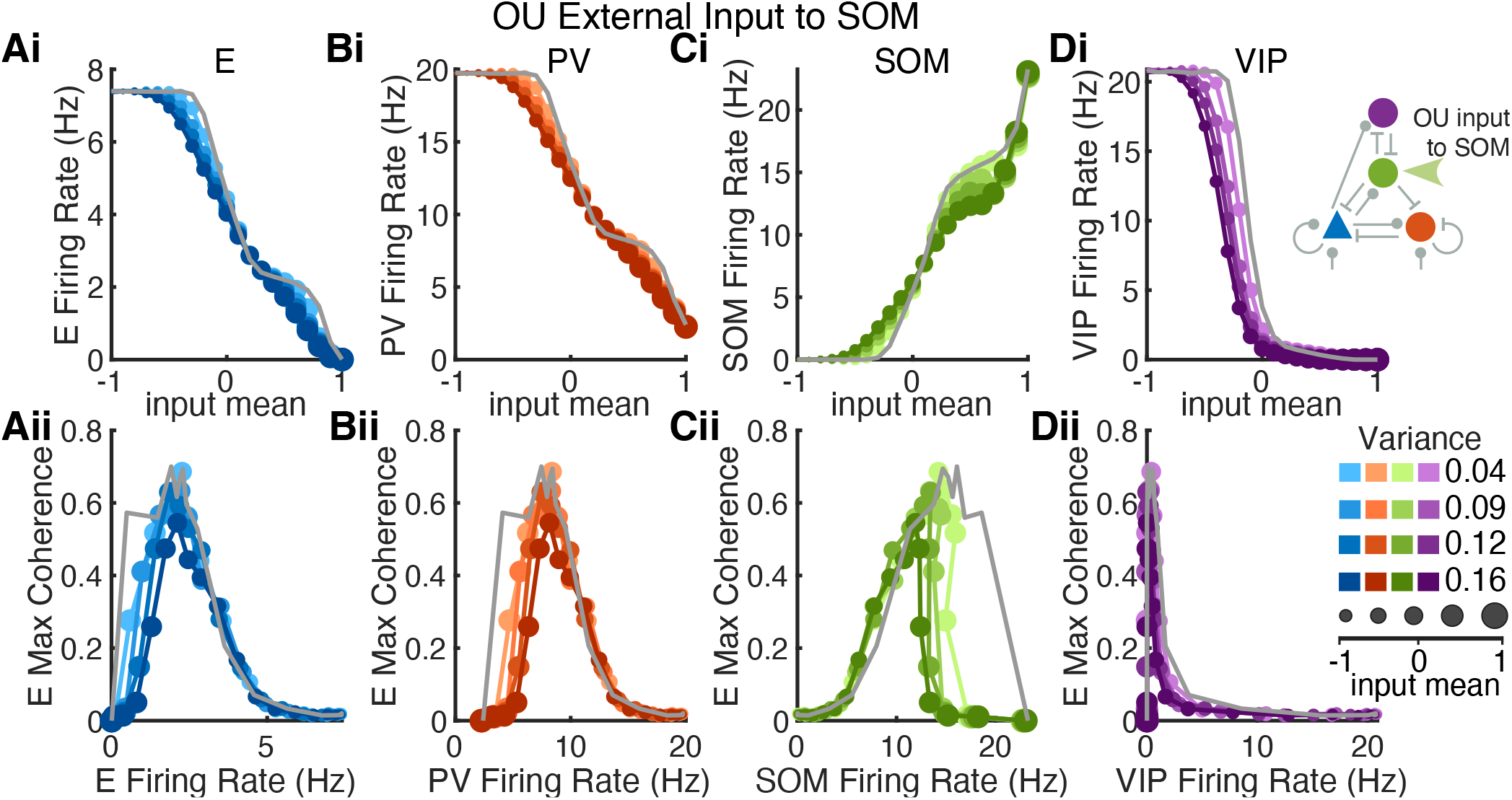
Weak impacts of dynamic noise input parameters on population rates and coherence. Related to Figure 8. Independent temporally-varying noise, modeled as an OU process of given mean (dot size) and variance (color shade), is applied to each SOM neuron. Columns show firing rate and coherence of (A) E, (B) PV, (C) SOM, (D) VIP populations. Row (i): Average firing rate of each cell population with respect to the mean value of OU input. Row (ii): The maximum coherence of E neurons compared to the population firing rates of each cell type. The grey curves are from the default network with SOM receiving static input.

**Figure S17:**
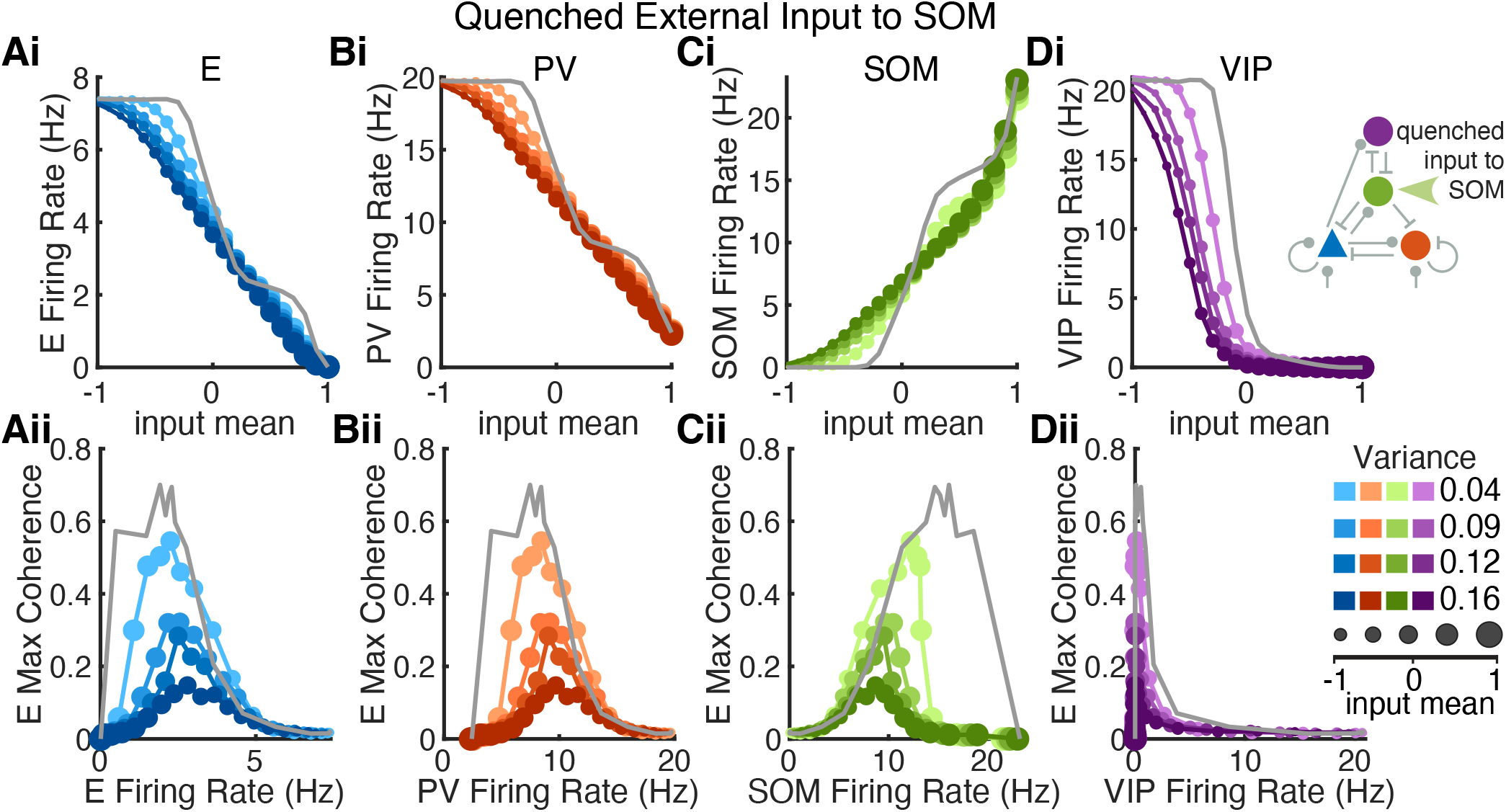
Impacts of quenched inputs on population rates and coherence. Related to Figure 8. Quenched input is spatially variable, but temporally invariant. Each SOM neuron receives an input value that is sampled from a Gaussian distribution with given mean (dot size) and variance (color shade). Columns show firing rate and coherence of (A) E, (B) PV, (C) SOM, (D) VIP populations. Row (i): Average firing rate of each cell population with respect to the mean value of the quenched input. Row (ii): The maximum coherence of E neurons compared to the population rates of each cell type. The grey curves are from the default network with SOM receiving static input.

**Caption for Movie S1**. Spiking activities of the spatially dependent spiking neuron network in the subcircuit asynchronous (SA) state (same parameters as in Figure 2A). Each dot indicates that the neuron at spatial position (*x, y*) fired within one millisecond of the time stamp shown on top. Color of each dot indicates the cell type of the neuron that fired (blue: E; red: PV; green: SOM; purple: VIP).

**Caption for Movie S2**. Same as Video 1 for the network in the weakly synchronous (WS) state (same parameters as in Figure 2B).

**Caption for Movie S3**. Same as Video 1 for the network in the strongly synchronous (SS) state (same parameters as in Figure 2C).

